# Single-cell analyses of axolotl forebrain organization, neurogenesis, and regeneration

**DOI:** 10.1101/2022.03.21.485045

**Authors:** Katharina Lust, Ashley Maynard, Tomás Gomes, Jonas Simon Fleck, J. Gray Camp, Elly M. Tanaka, Barbara Treutlein

## Abstract

Salamanders are important tetrapod models to study brain organization and regeneration, however the identity and evolutionary conservation of brain cell types is largely unknown. Here, we delineate cell populations in the axolotl telencephalon during homeostasis and regeneration, representing the first single-cell genomic and spatial profiling of an anamniote tetrapod brain. We identify glutamatergic neurons with similarities to amniote neurons of hippocampus, dorsal and lateral cortex, and conserved GABAergic neuron classes. We infer transcriptional dynamics and gene regulatory relationships of postembryonic, region-specific direct and indirect neurogenesis, and unravel conserved signatures. Following brain injury, ependymoglia activate an injury-specific state before reestablishing lost neuron populations and axonal connections. Together, our analyses yield key insights into the organization, evolution, and regeneration of a tetrapod nervous system.

---

Comparison of brains among animals has been a major means to analyze the evolutionary origin and diversification of this remarkable structure. Comprehensive single-cell (sc) and single-nucleus (sn) messenger RNA sequencing (RNA-seq) and spatial transcriptomics have greatly increased the resolution of cell identity and development of the vertebrate brain. Cell types from the dorsal region of the telencephalon, that in mammals includes hippocampus, amygdala, claustrum, olfactory (piriform) cortex and the highly elaborate neocortex, have been deeply resolved and compared between amniotes, including turtle, lizards, birds, mouse and human (1–6). These studies have revealed previously unappreciated evolutionary relationships in cell types and brain regions, such as the larger diversification of glutamatergic compared to GABAergic neurons, and the identification of the dorsal ventricular ridge as the direct sauropsid correspondent to the mammalian ventral pallium derivatives. With the aforementioned results deriving from the studies of amniote telencephalic cell types the question of conservation beyond amniotes arises. The salamander axolotl (*Ambystoma mexicanum*), as part of the amphibians, represents one of the closest living relatives to amniotes and is therefore suited to address this question. The fairly simple nature of the salamander brain makes it an important species for comparative studies of brain cell types, neuronal connectivity and function.

Comparative studies can also reveal how cellular processes and gene regulatory mechanisms have been adapted in distinctive animal traits such as regeneration. Notably, the axolotl is able to regenerate the telencephalon after removal of the dorsal region (7, 8). Regeneration of lost brain tissue occurs through proliferation of ependymoglia cells whose daughter cells differentiate into neurons over time, a process called neurogenesis. Neurogenesis is a universal vertebrate feature found in the developing brain and in certain adult brain niches (9–12). In the mouse, cells of the adult subventricular zone (SVZ) undergo continuous neurogenesis throughout life, however after brain injury neurogenesis is almost absent (13, 14). In aquatic animals such as axolotls that display long-term body growth, post-embryonic neurogenesis is observed in multiple brain regions. The molecular relationship between neurogenesis seen in salamanders and mammals has not been explored. Furthermore, similarities and differences between homeostatic and regenerative neurogenesis in the salamander brain are unclear.

Here, to understand the organizational features of the axolotl telencephalon, we employed snRNAseq and spatial transcriptomics. We analyzed the excitatory and inhibitory neuronal cell types, ependymoglia and neuroblasts present in different regions and defined their similarities to amniote telencephalic cell types. To delineate the cellular and molecular dynamics of homeostatic neurogenesis in the axolotl and its relation to adult neurogenesis in mice we utilized clonal tracing, trajectory analysis and multiomic sequencing. Analyzing regenerative neurotgenesis we determined the similarities and differences to homeostatic neurogenesis and found that regenerated neurons reestablish neuronal input from other regions of the telencephalon. Together, our comprehensive analyses of the axolotl telencephalon yielded key insights into the organization, evolution and regeneration of a tetrapod nervous system.

## Single-nucleus RNA-seq atlas of the axolotl telencephalon

We used single-nucleus RNA-sequencing to generate a comprehensive dataset of the cell types and states from the axolotl telencephalon. We micro-dissected the telencephalon into medial (containing medial pallium and septum), dorsal (containing dorsal pallium) and lateral (containing lateral pallium, ventral pallium and dorsal striatum) regions, and additionally profiled all these regions as a whole (Fig. 1A and fig. S1A). We computationally integrated these data, obtained through two different library chemistries. Clustering of 48,136 nuclei identified 95 molecularly distinct clusters including neuronal and non-neuronal cells (Fig. 1, B to E and fig. S1, B to E). We annotated major non-neuronal clusters as endothelial cells (*Col4a1+*), and glial cells including oligodendrocytes (*Musk+*), microglia (*Csf1r+*), and ependymoglia (*Gli2+*). Neuronal clusters included glutamatergic excitatory neurons (*Slc17a6/7+*), GABAergic inhibitory interneurons (*Gad1+, Gad2+*), and neuroblasts (*Mex3a+, Gli2-*) (fig. S1E). Each major cell type was present in each micro-dissected region (Fig. 1C), and we provide a set of marker genes for each of the identified clusters at this resolution (Fig. 1, D to E and table S1). We performed immunofluorescence stainings and RNA hybridization chain reactions (HCR) in situ to localize major cell types in the tissue based on marker expression (Fig. 1F). GFAP+ ependymoglia cells line the ventricle of every region, while *Mex3a+* neuroblasts are sparsely distributed along the ventricle. GABAergic neurons were sparsely distributed along the medial, dorsal and lateral region and clustered together densely in the striatum, while glutamatergic neurons were located in the medial, dorsal, lateral, and ventral pallium. We identified small populations of MBP+ oligodendrocytes and IBA1+ microglia cells distributed throughout all regions.

**Fig. 1.**
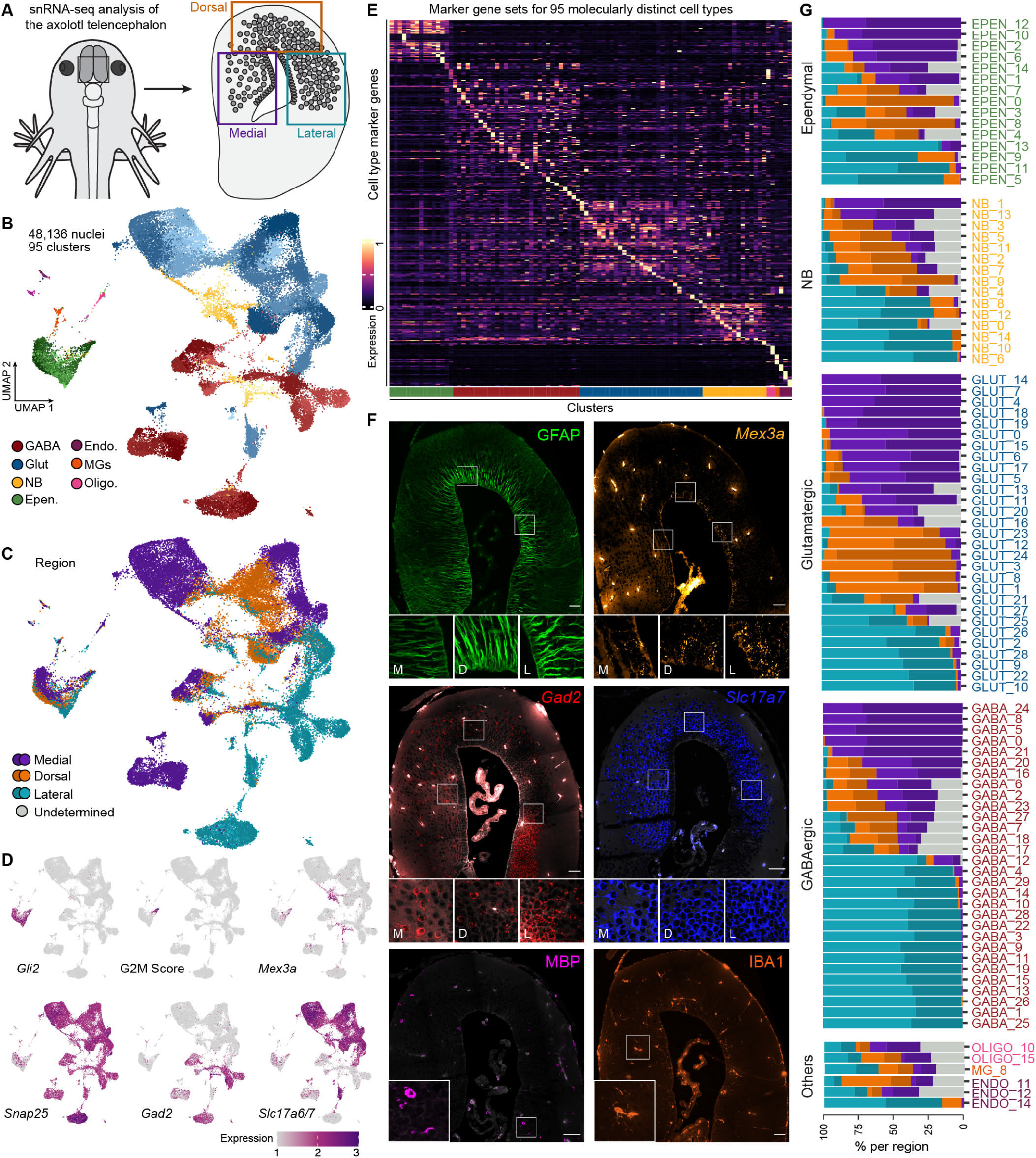
Single-nucleus RNA-sequencing reveals distinct ependymoglia, neuroblasts, and neurons in the medial, dorsal and lateral telencephalon. (A) Schematic highlighting the regions of the axolotl telencephalon used for single nucleus RNA sequencing. (B) UMAP of all cell types across all regions colored by cell-type class. Subtypes are shown in different shades. GABA, GABAergic neuron; Glut, Glutamatergic neuron; NB, neuroblast; Epen., ependymoglia cell; Endo., Endothelial cell, MGs, microglia; Oligo., oligodendrocyte. (C) UMAP plot of regional distribution of all cell types, shades indicate true region identity vs predicted regional identity. (D) UMAP plot of the expression of markers for ependymoglia, neuroblast and neuronal cell types. *Gli2* (ependymoglia cells), G2M score (active ependymoglia cells), *Mex3a* (neuroblasts), *Snap25* (neurons), *Gad2* (GABAergic neurons), *Slc17a6/7* (glutamatergic neurons) (E) Heatmap illustrates the expression of marker genes (table S1) for the 95 distinct cell types. (F) Antibody stainings and HCR in situ hybridizations for main cell types: GFAP (ependymoglia cells), *Mex3a* (neuroblasts), *Gad2* (GABAergic neurons), *Slc17a7* (glutamatergic neurons), MBP (oligodendrocytes), IBA1 (microglia cells). Scale bars are 100 µm. (G) Stacked barplots illustrating the regional distribution of the populations of cells. GABA, GABAergic neuron; Glut, Glutamatergic neuron; NB, neuroblast; Epen, ependymoglia cell; Endo., Endothelial cell, MGs, microglia; Oligo., oligodendrocyte.

We next analyzed the abundance of each cell cluster in the sampled pallial regions (Fig. 1G and fig. S1F). Oligodendrocytes, microglia and endothelial cell clusters originated from each region in similar proportions. In contrast, ependymoglia, neuroblast, glutamatergic and GABAergic neuron clusters each showed strongly differential region contributions, with some cell clusters being completely region-restricted, while others could be found throughout the sampled pallium regions. These data provide an overview of cell populations in the axolotl telencephalon, and suggest substantial regional specificity in neurogenic programs.

## Axolotl glutamatergic neuron types show similarities to amniote hippocampus, dorsal and lateral cortex

Glutamatergic neurons of the amniote pallium and cortex show a high degree of diversification and it has become clear in the last years that the molecular signatures of dorsal pallial pyramidal neurons are different across amniote species (1, 2). Until now, no single cell/nucleus dataset of a tetrapod anamniote species (salamanders or frogs) has been generated to understand how telencephalic glutamatergic neuron types relate to amniote glutamatergic neuron types. Our data revealed 29 glutamatergic neuron clusters that were distributed differentially in medial, dorsal and lateral regions (Fig. 2, A to C, and fig. S2A). To gain an understanding of potential homologies of axolotl glutamatergic subtypes to cortical cell types, we performed cluster correlation analysis of all axolotl glutamatergic neuron clusters to a turtle pallium dataset (2) and a mouse telencephalon dataset (3) (Fig. 2, D to E and fig S2, B to C). We made two separate comparisons using both species-shared differentially expressed genes and differentially expressed transcription factors (TFs) to potentially distinguish between convergent evolution versus homology, and focused on glutamatergic clusters that showed similarity with both gene sets as indicators of conservation.

**Fig. 2.**
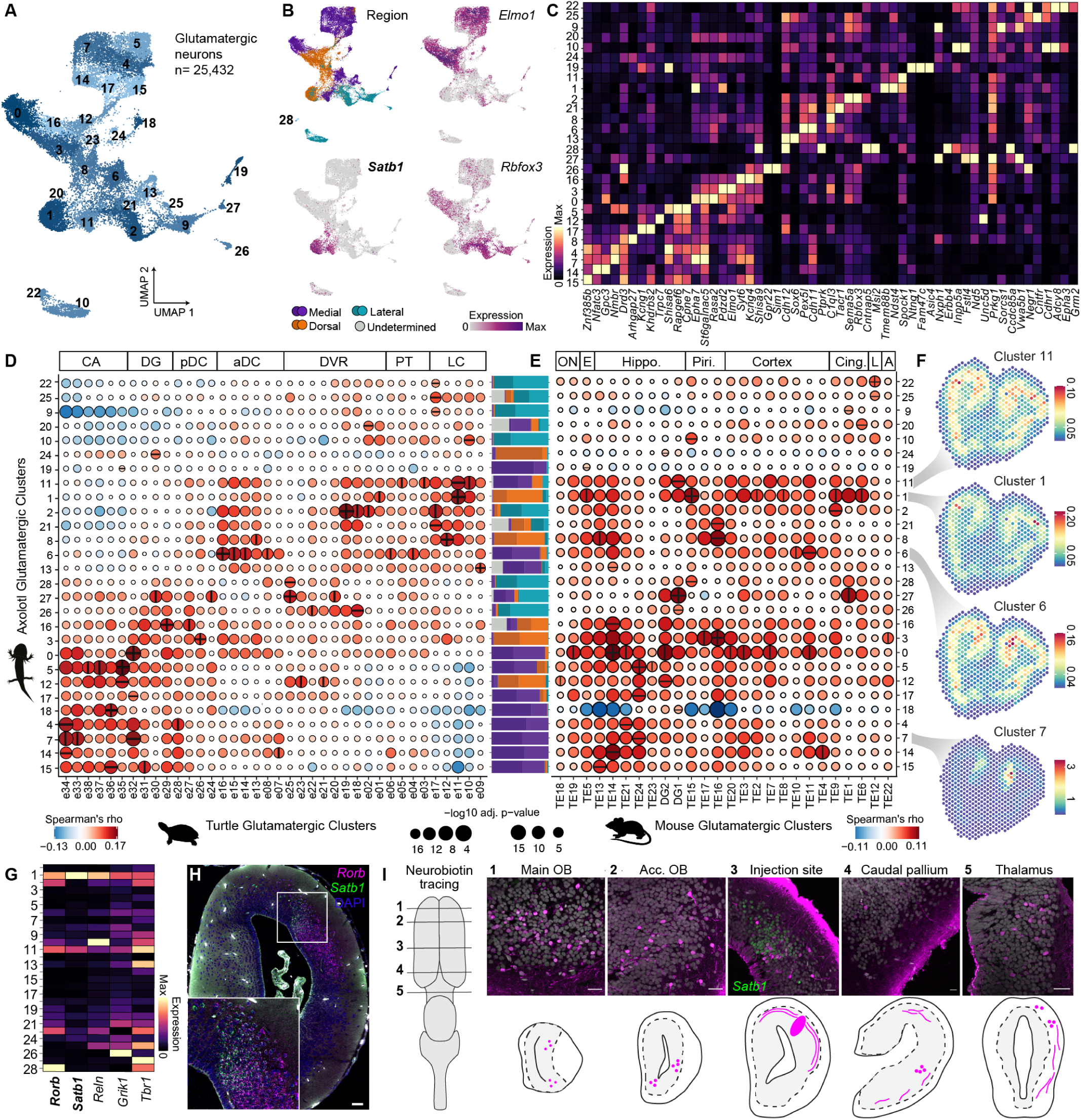
Glutamatergic neurons in the axolotl telencephalon show vertebrate-conserved signatures. (A) UMAP plot of 29 glutamatergic neuron types. (B) UMAP plots of the regional distribution of glutamatergic neurons types and the expression of three marker genes *Elmo1, Satb1* and *Rbfox3*. (C) Heatmap illustrating the expression of top markers for each glutamatergic neuron cluster. (D) Correlation analysis between expression profiles of axolotl glutamatergic neuron types and turtle glutamatergic neuron types (data from (2)). (E) Correlation analysis between expression profiles of axolotl glutamatergic neuron types and mouse glutamatergic neuron types (data from (3)). (F) Spatial mapping of glutamatergic neuron clusters 11, 1, 6, and 7. (G) HCR in situ hybridization for *Satb1* and *Rorb* (H) Heatmap illustrating the expression of *Satb1, Rorb, Reln, Grik1* and *Tbr1*. (I) Projection mapping of input into glutamatergic cluster 1 using Neurobiotin-mediated antero- and retrograde tracing. Dots indicate locations of cell bodies, lines locations of fibers.

Anatomical, developmental and functional evidence suggests that the amphibian medial pallium is homologous to the mammalian hippocampus (15, 16). In agreement with this we found that the majority of medial glutamatergic clusters (Glut 4, 5, 7, 15 and 18) as well as two dorsal clusters (Glut 12 and 16) matched either turtle Cornu Amonis (CA) or dentate gyrus (DG) clusters (Fig. 2D and fig. S2B). Correlations to the mouse dataset revealed a similar picture, medial glutamatergic clusters (Glut 5, 7, 14 and 17) matched hippocampus-related clusters (Fig. 2E and fig. S2C). Projection mapping of the frog medial pallium has led to the conclusion that subiculum and CA divisions but no DG exist (15). We found three clusters that showed one-to-one cluster correspondence to turtle CA1 (Glut 12), CA3 (Glut 4) and DG (Glut 16), however correlations to mouse did not reveal the same correspondences. To reveal the locations of hippocampus-matching glutamatergic clusters and unravel a potential subdivision into CA or DG we performed spatial transcriptomics (Fig. 2F, fig. S2D, and fig. S3, A to D). We found that all clusters except Glut 18 showed exclusive localization in the medial pallium, however clear subdivisions were not detectable. HCR for *Etv1, Prox1*, and *Lmo3* also failed to reveal clear subdivisions within the medial pallium (fig. S2E). These data show that neurons of the axolotl medial pallium have transcriptional similarities to amniote hippocampal neurons, but a clear distinction into CA1, CA3 and DG cannot be observed.

One cluster from the medial fraction (Glut 6) showed correlations to clusters of the turtle anterior dorsal cortex (aDC) and the pallial thickening (PT). The aDC has been previously reported to contain the most homologous cell types to mammalian cortical cell types (2). Interestingly, the correlation to mouse showed that Glut 6 also matched to clusters belonging to cortical cells (Fig. 2E). To define the location of Glut 6 we turned to our spatial transcriptome dataset (Fig. 2F). We found that this cluster was predicted to be located at the border between the medial and dorsal pallium.

One major function of the amphibian pallium is to process olfactory input (17, 18). In the sauropsid forebrain the lateral cortex (LC) is the main olfactory-input recipient region (19, 20). One axolotl glutamatergic cluster (Glut 1) showed strongest correlation to the turtle anterior LC (aLC) (Fig. 2D and fig. S2B). Neurons in the turtle aLC are considered homologous to neurons in the mammalian olfactory (piriform) cortex and interestingly Glut 1 also correlated to a mouse piriform cortex cluster (Fig. 2E and fig. S2C). The piriform cortex contains so-called semilunar cells, which receive strong excitatory inputs from the olfactory bulb (OB) and are characterized by the expression of *Rorb, Reln, Grik1, Tbr1, Fezf2* (21). We detected expression of *Rorb, Reln, Grik1* and *Tbr1* in Glut 1 (Fig. 2G). Moreover this cluster was characterized by strong expression of *Satb1* which we used in combination with *Rorb* expression to define its precise location which was consistent with the location of this cluster in the spatial transcriptomic dataset (Fig. 2, F and H). The mammalian piriform cortex receives input from the OB, entorhinal cortex, orbitofrontal cortex and the amygdala (22). We therefore analyzed the conservation on the level of connectivity by injecting the bidirectional neuronal tracer Neurobiotin into the Glut 1 region (Fig. 2I). Wholemount staining as well as sections allowed us to detect cell bodies in the main and accessory OB, regions of the caudal pallium likely corresponding to the lateral amygdala (23) as well as the thalamus, indicating that neurons in these regions project into Glut 1. This shows that the axolotl pallium contains a cell population with transcriptional similarity to amniote olfactory cortex neurons and the neuronal projections targeting this population suggest a conserved role in olfactory processing.

## Axolotl GABAergic neurons show conserved features with amniotes

We identified 30 diverse clusters (n = 15,665 cells) of GABAergic (*Gad1+*/*Gad2+*) neuronal cells (Fig. 3, A to B) in the axolotl. In many vertebrates, GABAergic interneurons are born in the three ganglionic eminences, lateral ganglionic eminence (LGE), caudal ganglionic eminence (CGE) and medial ganglionic eminence (MGE), and migrate to the pallium during development (24, 25). Comparative single cell transcriptomic analysis in mammals, lizards, turtles and songbirds have revealed that GABAergic interneurons are deeply conserved between amniote species and express shared sets of TFs (1, 2). To gain a better understanding of the conservation of axolotl GABAergic clusters we analyzed the expression of TFs known to define GABAergic cell classes and performed cluster correlation analysis to a turtle pallium dataset (2) (Fig. 3, C to D, and fig. S4A).

**Fig. 3.**
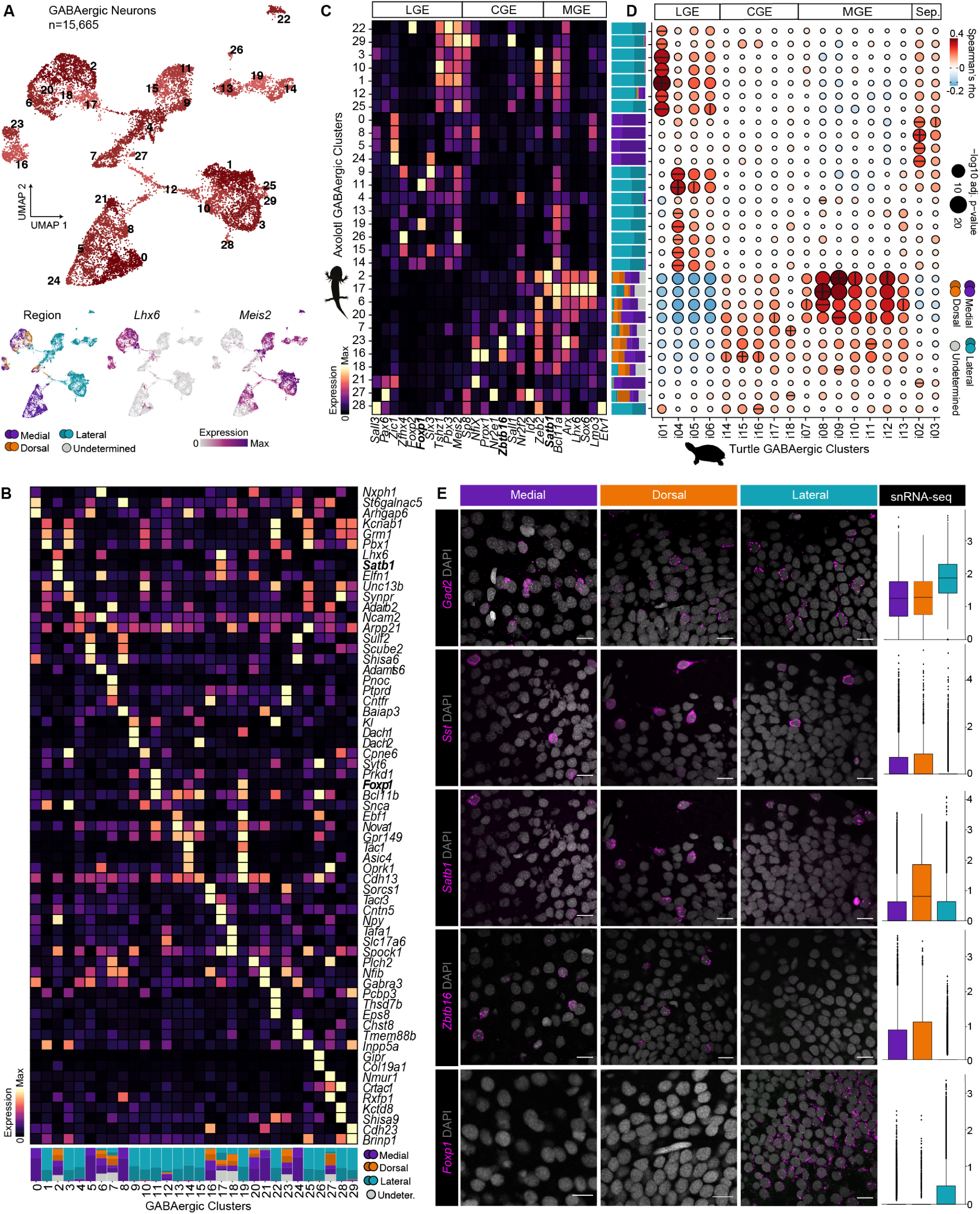
GABAergic neuron classes are conserved between axolotl and turtle. (A) UMAP plots of 30 GABAergic neuron types (top) and regional distribution of GABAergic neurons types as well as two marker genes *Lhx6* and *Meis2*. (B) Heatmap illustrating the expression of differential expressed markers for each GABAergic neuron cluster. (C) Heatmap illustrating the expression of transcription factors known to define GABAergic subtypes (LGE-, CGE-, MGE-derived) for each axolotl GABAergic neuron type. (D) Correlation analysis between expression profiles of axolotl GABAergic neuron types and turtle GABAergic neuron types (data from (2)). (E) HCR in situ hybridizations and snRNAseq quantifications for LGE-, CGE- and MGE-derived GABAergic neurons types (*Gad2*, pan; *Sst, Satb1*, MGE; *Zbtb16*, CGE, *Foxp1*, LGE). Scale bars are 25 µm.

Expression of conserved TFs allowed us to identify putative LGE-, CGE-, and MGE-derived (called LGE-, CGE-, and MGE-like from hereon) clusters in the axolotl dataset (Fig. 3C). Correlation analysis revealed that 13 out of 14 LGE-like clusters (GABA 1, 3, 9-15, 19, 22, 25 and 26) showed similarities to turtle LGE-derived clusters. Moreover, 2 out of 4 CGE-like clusters (GABA 7 and 16) and 5 out of 7

MGE-like clusters (GABA 2, 6, 17, 20 and 23) showed significant correlations to turtle CGE-derived and MGE-derived clusters respectively. Our analyses additionally identified 5 axolotl clusters (GABA 0, 5, 8, 21 and 24) composed of medial cells likely derived from the septum, which showed strong correlations to the turtle GABAergic neurons assigned to septum.

Turtle LGE-, CGE- and MGE-derived GABAergic neurons could be further subdivided into different GABAergic classes (2) but their existence in the axolotl was unknown. We found that 11 out of 13 axolotl LGE-like clusters correlated with either turtle striatum or olfactory bulb GABAergic cells and the majority of these (8 clusters) also correlated to mouse striatum or olfactory bulb GABAergic cells (fig. S4, B to C). In contrast to that, only 1 out of 4 CGE-type clusters (GABA 16) correlated to turtle HTR3A VIP-like neurons. Surprisingly, all 7 axolotl MGE-like neuron clusters matched to turtle PV-like neurons by TF expression but had similarities to turtle SST neurons at the level of effector genes, suggesting evolutionary divergence of these cells.

In mammals and turtles, MGE- and CGE-derived interneurons are differently distributed across cortical layers (2, 26). In axolotl, we found MGE-like neurons (*Sst+, Satb1+*) in all regions of the medial pallium. In contrast, these same neurons were enriched in the outer regions of the dorsal and lateral pallium (Fig. 3E and fig. S4D). CGE-like neurons (*Zbtb16+*) were located in all areas of the medial region but closer to the ventricle in the dorsal pallium. LGE-like striatal GABAergic neurons (*Foxp1+*) were found exclusively in the striatal region (fig. S4D). Finally, we spatially mapped LGE-, CGE- and MGE-like clusters and detected the majority of CGE- and MGE-like clusters located in all regions of the pallium as well as the striatum (fig. S4E). These data strongly suggest cell migration from putative CGE and MGE regions, whereas LGE clusters are predominantly located in the striatum, as in amniotes.

Together, these data show that the axolotl telencephalon contains putative LGE-, CGE- and MGE-derived as well as septal GABAergic neurons. While LGE-like striatum and olfactory bulb GABAergic classes showed strong transcriptional similarities between axolotl, turtle, and mouse, CGE- and MGE-like classes showed weaker and inconsistent correlations between analyses focusing on TFs alone or all differentially expressed genes.

## Ependymoglia cells in the axolotl telencephalon are regionally distinct

The predominant glial cells in the salamander central nervous system are ependymoglia, which are the source of new neurons in development, homeostatic growth, and regeneration in axolotl and newts (7, 8, 17, 27). The diversity of ependymoglia has only been addressed in the adult red-spotted newt telencephalon, where two ependymoglia types have been identified: quiescent type-1 cells that act as long-term stem cells and proliferative type-2 cells that are progenitor-like (28).

To examine the diversity of axolotl telencephalic ependymoglia, we subsetted the ependymoglia populations (3,590 cells) and identified 15 transcriptionally distinct cell clusters (Fig. 4A and fig. S5A). We grouped these clusters into three main types of ependymoglia that were present in all dissected regions: quiescent ependymoglia, active ependymoglia, and a novel ependymoglia type which we termed pro-neuro ependymoglia (Fig. 4A). Quiescent ependymoglia were characterized by expression of *Edn3*, active ependymoglia were characterized by expression of *Notch1* and a high score for G2M phase of the cell cycle and pro-neuro ependymoglia showed expression of neuron-related genes, such as a high expression of *Grin1* (Fig. 4, B to C, and fig. S5, B to D).

**Fig. 4.**
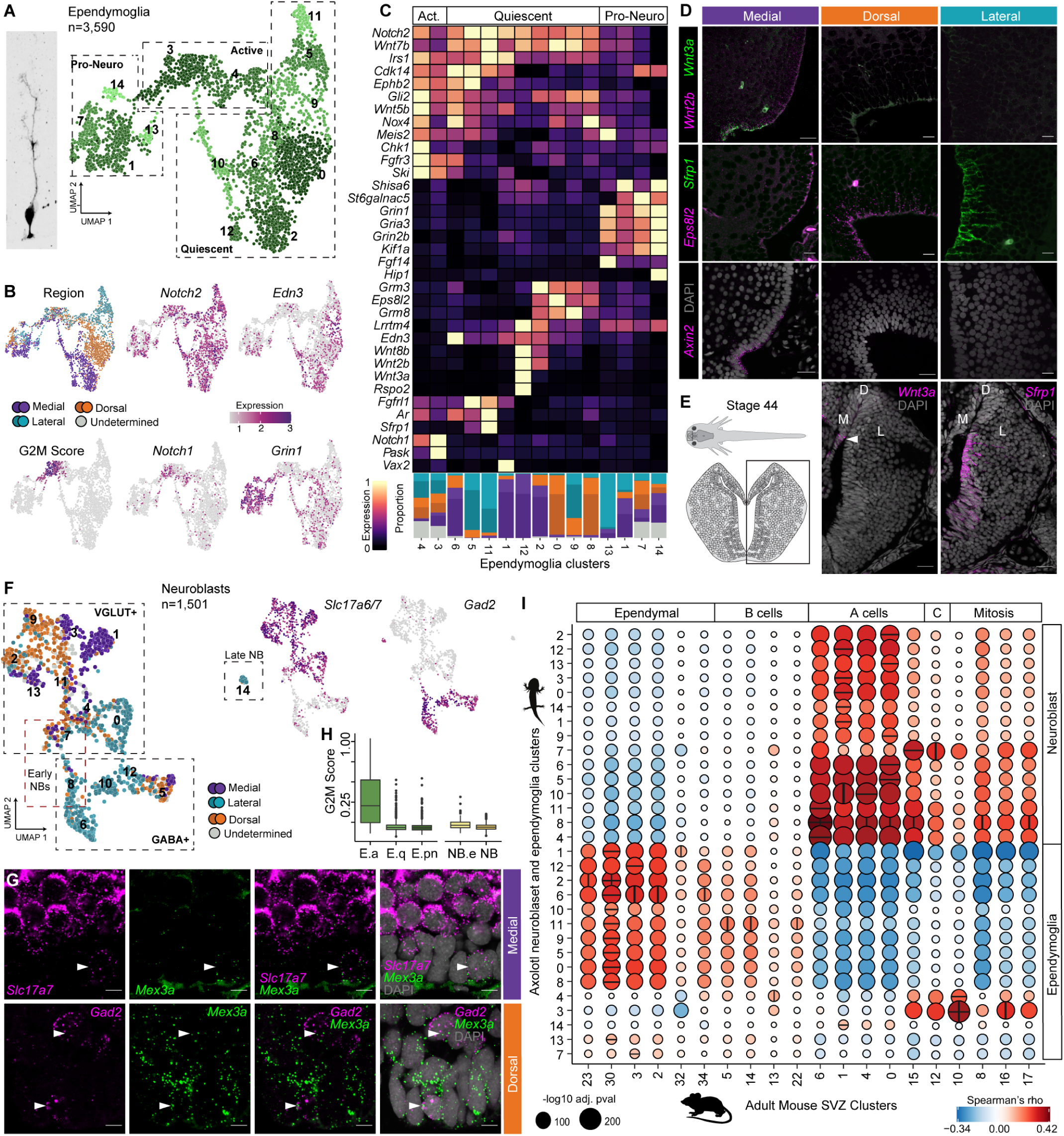
Ependymoglia and neuroblasts in the axolotl telencephalon are diverse and have regional signatures. (A) Morphology of an axolotl ependymoglia cell (left) and UMAP plot (right) of 15 ependymoglia clusters, boxes outline the three main cell types: quiescent ependymoglia, active ependymoglia and pro-neuro ependymoglia. (B) UMAP plot of the regional distribution of ependymoglia and UMAP plots colored by expression for *Notch2* (quiescent and active ependymoglia), *Edn3* (quiescent ependymoglia), G2M score (active ependymoglia), *Notch1* (active ependymoglia) and *Grin1* (pro-neuro ependymoglia). (C) Heatmap illustrating the expression of differentially expressed genes in quiescent, active, and pro-neuro ependymoglia. (D) HCR in situ hybridizations and snRNAseq quantifications for *Wnt2b, Wnt3a, Eps8l2* and *Sfrp1* in medial, dorsal and lateral regions. EdU staining after 2 consecutive injections within 2 weeks and snRNAseq qualifications for G2M and S phase score in medial, dorsal and lateral regions. Scale bars are 25 µm. (E) HCR in situ hybridizations *Wnt3a* and *Sfrp1* in the stage 44 developing pallium. Scale bars are 50 µm. (F) UMAP plots of the regional distribution 15 neuroblast clusters, black boxes outline the four main cell types: early neuroblasts, VGLUT+ and GABA+ neuroblasts, as well as late neuroblasts. Red box outlines the early neuroblast population. (G) HCR in situ hybridizations for *Scl17a7, Gad2*, and *Mex3a*. Arrows highlight co-expressing cells. Scale bars are 10 µm. (H) Boxplot of G2M score for quiescent, active and pro-neuro ependymoglia cells as well as neuroblasts. (I) Correlation analysis between expression profiles of axolotl neuroblasts, ependymoglia and adult mouse SVZ cell types (data from (37)).

In the mouse subventricular zone (SVZ), neural stem cell regional identities are maintained until adulthood and these identities underlie the types of neurons that can be generated (29–31). Strikingly, we found a clear distinction between medial-, dorsal- and lateral-derived ependymoglia (Fig. 4, C to D, and fig. S5E). These regional differences in gene expression were most notable for signaling pathway components (Fig. 4C). We detected strong expression of *Wnt8b* in a subset of medial quiescent ependymoglia, a gene known to be expressed in the medial region of the developing pallium in human, mouse and chicken and involved in patterning (32–34). In line with *Wnt8b* expression, we detected strong expression of *Wnt2b, Wnt3a* in medial ependymoglia. On the lateral side, cluster 11 (lateral ependymoglia) showed strong expression of *Sfrp1*, a gene known to be expressed in the antihem, a ventral pallial region in amniotes (35, 36). Dorsal ependymoglia were enriched for expression of the gene *Eps8l2*. To further spatially resolve the expression patterns and validate our snRNA-seq data, we performed HCR for *Wnt2a, Wnt3a, Axin2, Sfrp1* and *Eps8l2. Wnt2b, Wnt3a* and *Axin2* were expressed in a restricted domain of the medial pallium at the border to the septum. In contrast, *Ep8sl2* was detected in the dorsal region of the pallium, while *Sfrp1* showed expression in the lateral and ventral region (Fig. 4D). Since Wnt and Sfrp genes have been implicated in patterning of the pallium during development we additionally performed HCR in embryonic axolotl brains (stage 44). We detected *Wnt3a* in a small population of ventricular cells of the developing medial pallium and *Sfrp1* in ventricular cells of the lateral and ventral developing pallium, however *Ep8sl2* was not detected in the dorsal pallium (Fig. 4E and fig. S5F). These data show the axolotl telencephalon contains three main ependymoglia types (quiescent, active, and pro-neuro) that subset into regionally distinct subtypes and, barring pro-neuro ependymoglia, continue to express pallial patterning genes in the postembryonic brain.

## Regionally distinct neuroblasts express glutamatergic or GABAergic neuron markers

In addition to ependymoglia cells, we identified cells that we termed neuroblasts due to the expression of *Mex3a*, an RNA binding protein expressed in proliferating neuroblasts in the Xenopus laevis central nervous system (37). Additionally, neuroblasts were characterized by the absence of ependymoglia markers such as *Gli2, Aqp4* and *Kcnj10*. We detected 15 *Mex3a*-positive neuroblast clusters that could be divided into two distinct groups based on the expression of *Slc17a6/7* (10 clusters, VGLUT+) or *Gad1/2* (5 clusters, GABA+) (Fig. 4F and fig. S5, G to H). While VGLUT+ neuroblasts were predominantly present in medial and dorsal fractions with the exception of cluster 4, GABA+ neuroblasts were mostly found in the lateral dataset, with the exception of cluster 5. To validate the location of VGLUT+ or GABA+ neuroblasts in the ventricular region we performed HCR for Scl17a6 or Gad2 in combination with Mex3a (Fig. 4G). VGLUT+ neuroblasts were detected at the ventricle in all regions of the pallium while GABA+ neuroblasts were predominantly present at the ventricle in the striatum, with the exception of a few that could also be found in the pallium (Fig. 4G, fig. S5I). This pattern was also verified when mapping these clusters to our Visium data, where VGLUT+ and GABA+ neuroblasts were enriched in the same regions (fig. S5J) In contrast to intermediate progenitors found in other vertebrate brains, *Mex3a*-positive neuroblasts largely do not appear proliferative, based on a G2M score similar to quiescent and pro-neuro ependymoglia (Fig. 4H and fig. S5K). Two clusters of neuroblasts (clusters 7 and 8) showed an increased G2M score when compared to the rest and were therefore termed early neuroblasts.

To define the transcriptional similarities of axolotl ependymoglia and neuroblasts to neural stem and progenitor cells in other species, we performed cluster correlation analysis to an adult mouse SVZ dataset (38) (Fig. 4I and fig. S6A) and a mouse developmental cortex dataset (39) (fig. S6, B and C). Comparison of axolotl ependymoglia to mouse cells revealed that quiescent ependymoglia correlated either with mouse ependymal cells (adult and development) or B cells (adult), whereas active ependymoglia (clusters 3 and 4) showed strong correlation to mitotic cells (adult) including dividing A cells (cluster 16) or developmental apical and intermediate progenitors. Interestingly, active ependymoglia cluster 4 also showed correlations to B cell cluster 13, which has been described as the activated B cell population. Among the pro-neuro ependymoglia only cluster 1 showed strong correlation to ependymal cells (adult and development), whereas clusters 7 and 13 weakly correlated to ependymal cells (adult) or migrating neurons (development) and cluster 14 weakly correlated to A cells.

Comparison of axolotl neuroblasts to the adult mouse SVZ dataset revealed strong correlation to A cells which was supported by the strong correlation to migrating neurons in the developmental dataset (fig. S6, B and C). Interestingly, we found that early neuroblast clusters 7 and 8 correlated also to either C cells or mitotic cells. Furthermore, these two clusters showed strongest correlation to A cell clusters 6 and 15, which were defined previously as dividing neuroblasts/early A cells, as well as to intermediate progenitor cell clusters in the developmental dataset (Fig. 4I and fig. S6A). Together, these results show that the axolotl telencephalon contains neuroblast populations that already express neurotransmitter signatures of downstream neurons. Moreover, neuroblasts are most similar to mouse progenitor cells while ependymoglia harbor transcriptional similarities to mouse ependymal cells as well as neural stem cells.

## Transcriptional dynamics of postembryonic glutamatergic neurogenesis

BrdU incorporation assays have shown that neurogenesis in the telencephalon continues during postembryonic life of the axolotl (7). We used a Cre-loxP-mediated tracing approach labeling ependymoglia to address whether they behave as self-renewing stem cells and to determine the clonal patterns they generate during post-embryonic neurogenesis (Fig. 5A). Using this approach, we uncovered distinct neurogenesis patterns in medial, dorsal and lateral regions. Medial and dorsal domains contained continuous clones, which spanned from the ventricular zone up into the neuronal layers, indicating a tree-like growth mode (stacking) reminiscent of postembryonic neurogenesis in the zebrafish pallium (40). In contrast, the clonal patterns in the lateral domain were fragmented, with labeled neurons separated from ependymoglia indicating neuronal migration.

**Fig. 5.**
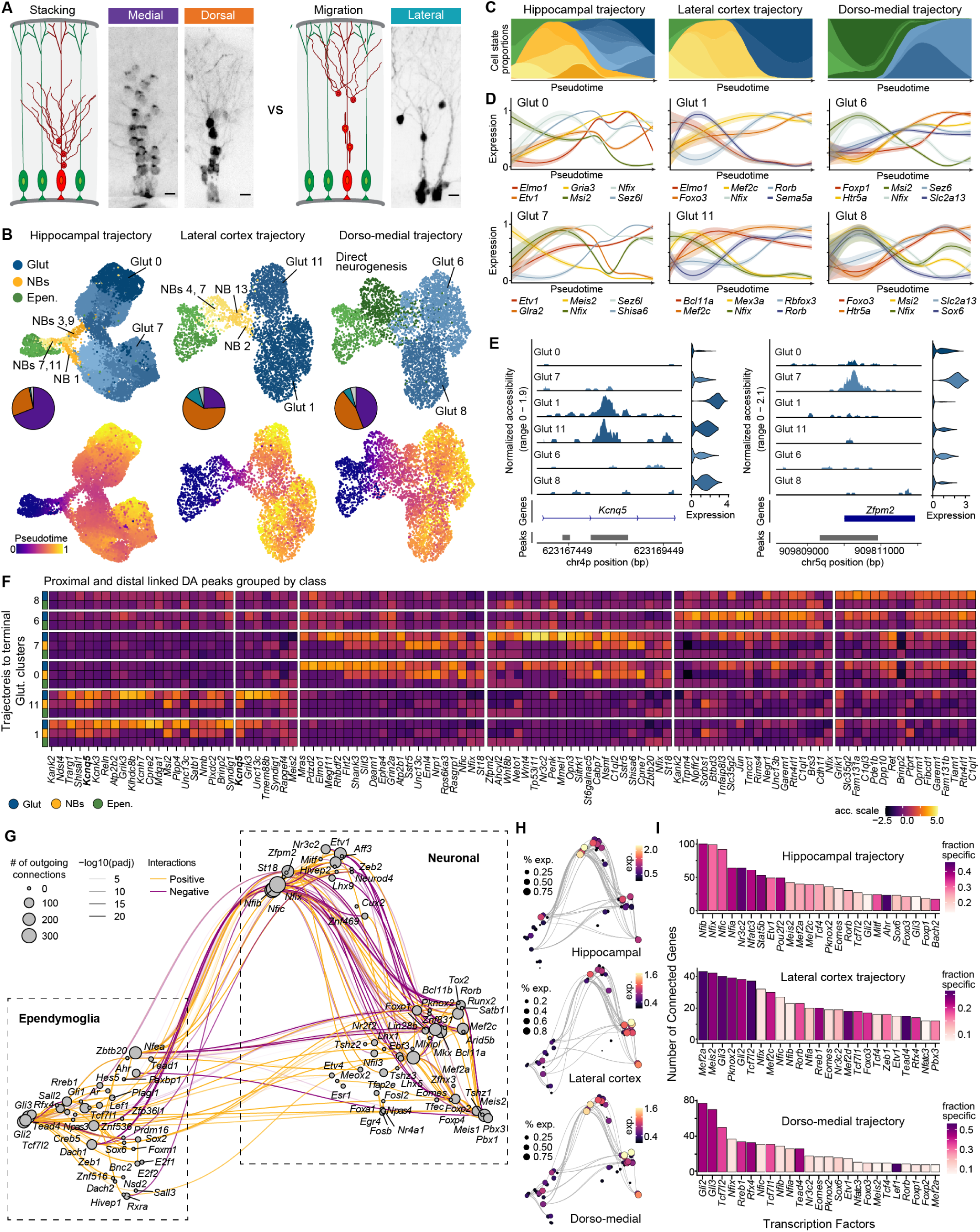
Distinct gene regulatory programs underlie adult neurogenesis in the different regions of the axolotl telencephalon. (A) Schematic describing the outcomes of Cre-loxP fate mapping performed to assess clonal dynamics and potential clone shapes during homeostatic neurogenesis of the axolotl telencephalon adjacent to measured clonal patterns in medial, dorsal and lateral pallium. (B) Glutamatergic trajectories reflecting adult neurogenesis of axolotl neurons matched to amniote hippocampus, lateral cortex and the dorso-medial pallium. UMAPs colored by cell types (top) and pseudotime (bottom). Pie charts represent the regional composition of neuron clusters. (C) Pseudotemporal cell type progression from ependymoglia to glutamatergic neurons during neurogenesis in the three trajectory groups. (D) Pseudotemporal gene expression changes during neurogenesis for each terminal branch from the trajectories of each group. (E) Representative example peaks associated with *Kcnq5* (left) and *Zfpm2* (right). (F) Heatmap of chromatin accessibility changes in distal and proximal elements for glutamatergic clusters 8, 6, 7, 0, 11, and 1. (G) UMAP embedding of the inferred gene modules based on co-expression and inferred interaction strength between transcription factors. Size represents the number of connections for each transcription factor. (H) Trimmed GRN UMAP embedding of the inferred gene modules based on co-expression and inferred interaction strength between transcription factors for Hippocampal (top), Lateral cortex (middle), and Dorsal-medial (bottom) trajectories. Color scale indicates expression and size represents the percent of cells expressing. (I) Barplot of the top 25 transcription factors ranked by number of connections for each transcription factor.

We used RNA velocity based trajectory inference (41, 42) to explore the cellular and molecular dynamics of postembryonic neurogenesis. We focused our analysis on glutamatergic neurons, since these are known to be locally generated, whereas GABAergic neurons migrate across the pallium from the striatum and thus our data likely misses some corresponding progenitor populations (24, 25). We focused on transitions from active ependymoglia to the most differentiated glutamatergic neurons. To construct trajectories we first identified the key VGLUT-positive neuroblast populations that have highest transcriptional similarity to the respective glutamatergic neuronal clusters (fig. S7A) and are therefore able to connect active ependymoglia to neuronal clusters in a trajectory. Using these groups of clusters we then constructed five trajectories representing different region-specific neurogenesis (Fig. 5B and fig. S7B). While all trajectories were rooted at the active ependymoglia, not all trajectories utilized neuroblast intermediates (Fig. 5, B to C and fig. S7B). Specifically, hippocampal neuronal clusters (Glut 0, 12, 14, 15, 16, 17, 3, 4, 5, and 7), lateral cortex clusters (Glut 1 and 11), and laterally derived clusters including the lateral pallium group (Glut 2, 9, 21, 25) and the *Eomes* group (Glut 10, 22) all utilized neuroblast intermediates. In contrast, the dorso-medial neuronal clusters (Glut 6, 8, and 20) formed a group with markedly lower correlations with neuroblast clusters 7, 11, and 4 (the least differentiated), and were thus inferred to originate through direct neurogenesis using pro-neuro ependymoglia (clusters 1, 14, and 7) as intermediates. For each trajectory we identified genes with varying expression along pseudotime revealing many trajectory specific genes, but also some genes with consistent pseudotemporal expression across trajectories regardless of their region specificity (Fig. 5D).

To resolve the gene regulatory relationships underlying glutamatergic neurogenesis, we leveraged our single-nucleus multiome-sequencing of the axolotl whole pallium. We identified proximal and distal candidate regulatory regions differentially accessible in each of the terminal glutamatergic neuron clusters and assessed at which stage in the respective trajectory they become accessible (Fig. 5, E to F, fig. S7, C to E, and table S2). Notably, most elements identified as specific for a given terminal glutamatergic cluster already obtained accessibility in the corresponding neuroblast clusters earlier in the trajectory.

We inferred a gene regulatory network (GRN) using Pando (43), by combining gene expression and chromatin accessibility measurements. A UMAP embedding of this GRN revealed distinct groups of TFs, corresponding to the transition from ependymoglia to glutamatergic neurons (Fig. 5G). To better understand how gene regulation differs between neuronal trajectories, we performed differential accessibility analysis and identified regulatory regions enriched within each trajectory. Based on these regions, we constructed subnetworks of the GRN reflecting trajectory-specific regulatory features (Fig. 5H and fig. S7F). This allowed us to identify the TFs with highest centrality in each trajectory, such as *Nr3c2* in hippocampus, *Rorb* in lateral cortex, *Tead4* in dorso-medial cortex, as well as *Tcf7l2* in lateral and dorsomedial cortex (Fig. 5I and fig. S7G). Interestingly, *Nfix* was one of the most central TFs in all subnetworks, however the regulomes controlled by *Nfix* were distinct for each respective trajectory (fig. S7, H to I). Together these data highlight the regulatory relationships shaping neuronal diversification in the axolotl telencephalon.

## Ependymoglia activate an injury-specific state before the start of neurogenesis during regeneration

To study the cellular and molecular dynamics during axolotl pallium regeneration, we implemented Div-Seq (44), which combines snRNAseq with EdU labeling of S-phase cells, thereby labeling proliferating cells and their progeny. We injured the dorso-lateral region of the pallium (including the *Satb1+, Rorb+* domain) by excising a 1mm by 1mm region and applied Div-Seq throughout regeneration by labeling cells with EdU at 2, 5, 12, 19 and 26 days post injury (dpi) and collecting EdU+ cells at 1, 2, 4, 6, 8 and 12 weeks post injury (wpi) (Fig. 6A).

**Fig. 6.**
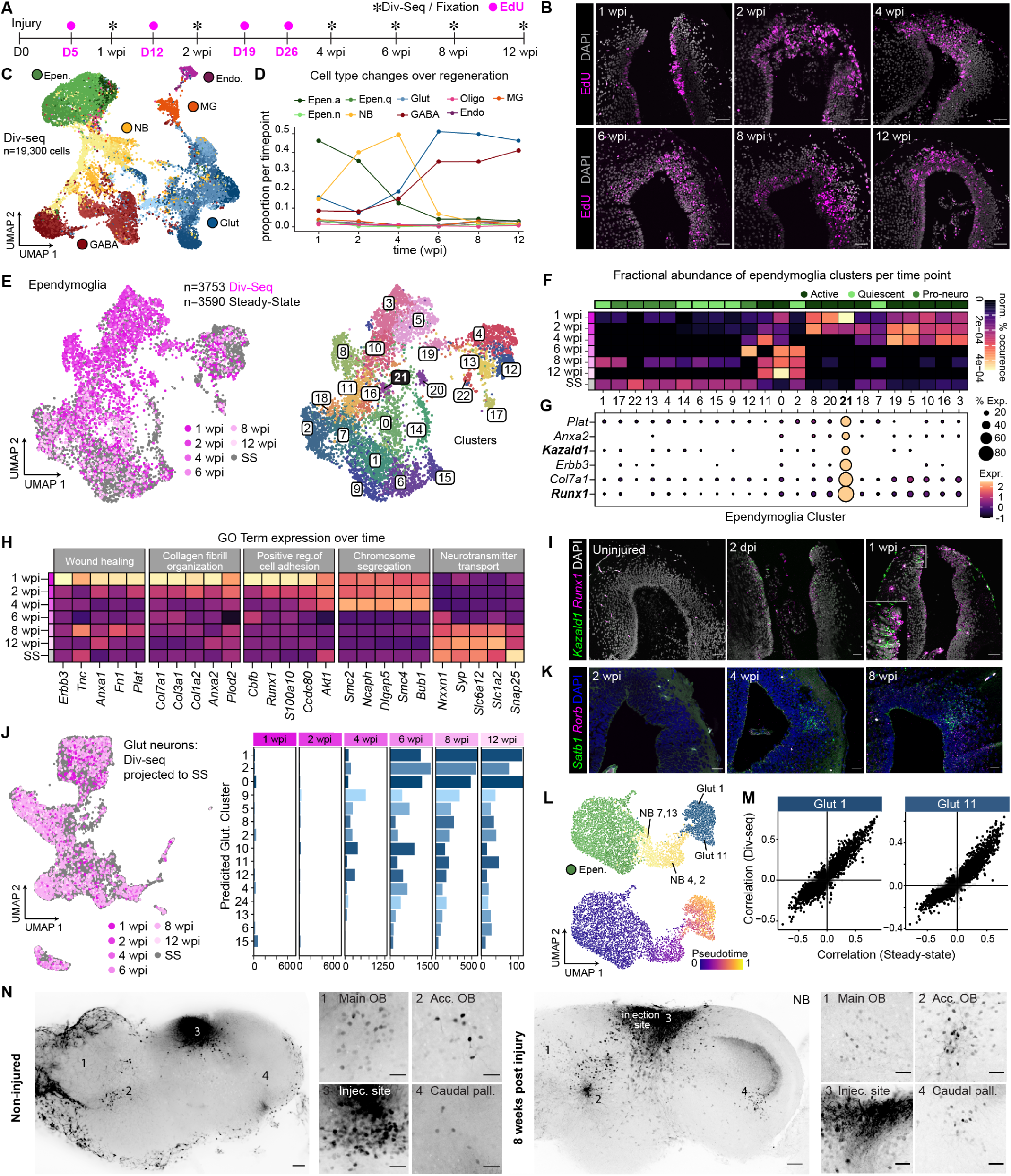
Ependymoglia progresses through an injury-specific state prior to regeneration of lost neuronal populations and connection patterns. (A) Schematic describing the Div-Seq protocol used during axolotl pallium regeneration. (B) EdU stainings of all regeneration time points. Scale bars are 50 µm. (C) UMAP plot of all EdU+ cells across all regeneration time points colored by cell-type class. Predicted cell clusters are shown in different shades. GABA, GABAergic neuron; Glut, Glutamatergic neuron; NB, neuroblast; Epen., ependymoglia cell; Endo., Endothelial cell, MGs, microglia. (D) Change in cell type relative abundance along regeneration time points. (E) Left: UMAP plot of all Div-Seq ependymoglia (shades of pink indicate different time points) and non-injured brain steady state (SS) ependymoglia (gray). Right: UMAP plot of clustering of all ependymoglia. (F) Heatmap of normalized cluster occurrence per time point. (G) Dotplot of selected ependymoglia cluster 21 DE genes. (H) Gene average expression change per time point, grouped by GO terms. (I) HCR in situ hybridization for *Kazald1* and *Runx1* in steady state, 2dpi and 1wpi. Scale bars are 50 µm. (J) Left: Correlation projection of all Div-Seq glutamatergic neurons (pink) to steady state glutamatergic neurons (gray). Right: Barplots of largest glutamatergic neuron clusters recovered throughout regeneration time points. (K) HCR in situ hybridizations for *Satb1* and *Rorb* throughout regeneration. Scale bars are 50 µm. (L) Trajectories reflecting regenerative neurogenesis of glutamatergic neuron clusters 1 and 11. UMAP colored by cell type (left) and pseudotime (right). (M) Correlations of lineage driver genes with the assignment probability for Glut 1 (top) and Glut 11 (bottom) trajectories, in regenerative (y-axis) and Steady-state (x-axis) neurogenesis. (N) Wholemount cleared neurobiotin stainings on non-injured (left) and 8 weeks post injury brains (right). Neurobiotin was injected in the *Satb1+, Rorb+* domain. Insets show labeled cell bodies in the olfactory bulb, accessory olfactory bulb, injection site and caudal pallium. Scale bars are 100 µm (overviews) and 50 µm (zooms).

To visualize the location of EdU+ cells during the regeneration process we fluorescently labeled EdU using click chemistry on regenerating brains (Fig. 6B). At 1 wpi the injury site was still open and EdU+ ependymoglia were present in medial and lateral wound-adjacent regions. At 2 wpi the injury site was starting to close by accumulation of EdU+ cells. Throughout the following time points, EdU+ cells remained accumulated at the regeneration site until tissue architecture was largely re-established.

Next, we investigated the transcriptomes of EdU+ cells during the regeneration time course. We used our steady-state data as a reference and identified all major cell types including ependymoglia, neuroblasts, glutamatergic and GABAergic neurons as well as endothelial cells and microglia (Fig. 6C). Interestingly, each cell type was represented in different proportions throughout the time course of regeneration (Fig. 6D and fig. S8A) reflecting a wave of neurogenesis induced by the injury. While at 1 wpi active ependymoglia constituted the majority of EdU+ cells, neuroblasts were the most abundant cell type at 2 and 4 wpi. Starting from 6 wpi, most of the EdU+ cells were glutamatergic and GABAergic neurons. In line with the results from the Div-Seq data, HCR and antibody staining for ependymoglia (GFAP+, *Eps8l2+, Sfrp1+*) and neuroblast (*Mex3a+*) populations, demonstrated that ependymoglia were recovering radial morphology and regional identity, while neuroblasts were accumulating at the wound site between 2 wpi and 4wpi (fig. S8B). We next inferred regional identities of Div-seq cells by transferring region labels from our steady-state data and found that early cell types (ependymoglia and neuroblasts) largely consisted of dorsal regional transcriptional identities. In contrast, EdU-labeled glutamatergic neuronal populations were predicted to have more mixed regional transcriptional identities and GABAergic neuronal populations were dominated by a lateral identity (fig. S8, C to D). This suggests that our Div-seq data largely captured cells in the acute injury area of the dorso-lateral pallium but also some cells derived from homeostatic neurogenesis in areas not associated with the injury site.

Previous studies on axolotl spinal cord regeneration showed that injury-induced ependymoglia activate a transcriptional signature similar to embryonic neuroepithelial cells (45). To understand whether pallium injury induces injury-specific changes in ependymoglia transcriptomes we compared ependymoglia from all regeneration time points to steady-state ependymoglia by clustering the harmonized datasets and assessing differential abundance and expression in each cluster across the time points (Fig. 6E). We found one cluster (cluster 21) strongly enriched at 1wpi that was absent from later time points and steady state (Fig. 6F). Notably, cluster 21 cells differentially expressed key genes such as *Kazald1* and were enriched for GO terms relating to wound healing and cell adhesion, indicating early response programs to injury (Fig. 6, F to H). Staining for *Kazald1* and *Runx1* confirmed absence of expression in the uninjured pallium and strong expression in a subpopulation of cells at 1wpi, with an induction of expression as early as 2 dpi (Fig. 6I).

Projection of Div-seq neuroblasts and neurons to the steady state data and classification based on label transfer showed that a majority of steady state populations had been re-established during regeneration (Fig. 6J and fig. S8, A and E to F). Among glutamatergic neurons, all but one steady state cluster was captured in the Div-seq data (fig. S8A), with the most expanded clusters predicted to be of dorso-medial origin (fig. S8, B to C). HCR staining for *Satb1+* and *Rorb+* demonstrated the recovery of *Satb1+, Rorb+* glutamatergic neurons starting at 4wpi and the re-establishment of a domain as had been observed in the uninjured tissue by 8wpi (Fig. 6K). We explored the dynamics of regenerative glutamatergic neurogenesis using RNA velocity-based trajectory analysis. We first determined the correlations of regenerating neuroblasts to regenerating glutamatergic neurons and found a strikingly similar correlation pattern as in homeostatic neurogenesis (fig. S8G). We then constructed trajectories of regenerative neurogenesis that were found highly similar to the trajectories of adult homeostatic neurogenesis, with similar driver gene correlations (Fig. 6, L to M, and fig. S8, H to K). Finally, we set out to determine whether the regenerated *Satb1+, Rorb+* neuron domain would re-establish afferent and efferent projections. We injected Neurobiotin into the *Satb1+, Rorb+* domain in non-injured as well as regenerated brains at 8 wpi and performed wholemount immunohistochemistry to identify cell bodies and projections. We found that, similar to non-injured brains, cell bodies were located in the olfactory bulb, accessory olfactory bulb and the caudal pallium (amygdala), indicating that the input from these regions is re-established in the regenerated pallium at 8 wpi (Fig. 6N).

## Discussion

Using single-nucleus RNA-, multiomic- and Div-sequencing along with spatial transcriptomics, Cre-loxP tracing, HCR, and antibody staining we have generated the first comprehensive single cell atlas of the axolotl telencephalon during homeostasis and regeneration. Comparative analysis with turtle and mouse datasets allowed us to reveal transcriptional similarities of axolotl telencephalon cell types facilitating an understanding of their conservation between tetrapods.

Axolotl glutamatergic neurons show similarity to amniote hippocampus, turtle aDC and olfactory cortex. Salamanders rely on olfaction during exploratory and feeding behaviors as well as stimulus response learning (17). We identified a glutamatergic neuron cluster (*Satb1+, Rorb+*) with strong transcriptional similarities to the turtle LC and mammalian olfactory cortex. The transcriptional profile we found in axolotl *Satb1+, Rorb+* cells is reminiscent of semilunar cells (mammals) or bowl cells (reptiles), which are excitatory cells receiving olfactory input (19, 21). Along with transcriptomic similarities, these axolotl neurons also showed input connectivity from the olfactory and accessory olfactory bulbs, indicating that their role in olfactory processing is conserved. To gain a better understanding of conservation it will be of importance to address the functional properties of these neurons and define their input and output connectivities in more detail at the level of synaptic connections. Our multiomic sequencing analysis has revealed differentially accessible regions in *Satb1+, Rorb+* glutamatergic neurons, which could be used in the future to achieve targeted expression of connectivity, optogenetic and chemogenetic tools.

We identified LGE-, CGE- and MGE-like as well as septal GABAergic neurons in the axolotl and found that they have transcriptional similarity to turtle and mouse GABAergic neurons. At the level of GABAergic classes we found a striking conservation of LGE-like striatal and olfactory bulb classes between axolotl and other tetrapods. In contrast to that, MGE-like and CGE-like classes were more divergent. The LGE and MGE have been found in all studied vertebrates including anamniotes such as lamprey, fish and amphibians through expression of conserved marker genes (46–49), but the existence of the CGE in anamniotes is unclear. The fact that we identified putative CGE-derived GABAergic neurons in our axolotl dataset hints at the existence of a CGE. GABAergic neuron migration has not been studied in salamanders until now and it will be important to determine the origin and timing of GABAergic neurogenesis in the putative ganglionic eminences during brain development. In axolotl, GABAergic neurogenesis likely continues in the postembryonic brain as we detected GABA+ neuroblasts in our dataset. While most GABA+ neuroblasts were derived from the lateral/striatum region we found one GABA+ neuroblast cluster (NB 5) containing cells from medial and dorsal pallium hinting at the possibility of local pallial GABAergic neurogenesis in the postembryonic axolotl brain, a phenomenon that has been observed in the developing primate brain (50, 51).

We also investigated the similarities of axolotl ependymoglia and neuroblasts to mammalian radial glia and progenitor cells. The mammalian SVZ contains ventricular ependymal cells acting as a supporting niche for radial glia-like B cells which are largely quiescent, but can self-renew or divide (12). We found that axolotl ependymoglia show transcriptomic signatures of both mouse SVZ ependymal cells and B cells. Functionally, axolotl ependymoglia act as long term stem cells during homeostatic neurogenesis. Interestingly, we uncovered that ependymoglia in different regions of the pallium maintain expression of developmental patterning genes such as Wnts, Fgfs and Sfrps even during postembryonic life. These secreted factors are thought to regulate dorsal and ventral pallial subdomain size during pallium development in different amniote species (52). It is tempting to speculate that due to the continuous neurogenic activity in the axolotl pallium expression of these factors might be responsible for maintaining the proportions of the respective pallial domains.

The mammalian SVZ also contains intermediate progenitor cells (IPCs) and migratory neuroblasts which express neuronal lineage markers depending on their GABAergic or glutamatergic fate (53, 54). In avian and mammalian brains IPCs are involved in indirect neurogenesis leading to the generation of a greater number of neurons and the enlargements of the dorsal cortex. In contrast, in reptiles cortical neurons are produced by direct neurogenesis (55). We identified different types of neuroblasts in the axolotl: early, potentially proliferating neuroblasts with high transcriptional similarity to mammalian IPCs, and late VGLUT+ or GABA+ neuroblasts with differential transcriptional similarities to mammalian neuroblasts. Importantly, the majority of axolotl VGLUT+ neuroblasts could be subdivided into different groups depending on their similarity to different glutamatergic neurons, implicating potential lineage restriction. However, one of the inferred neurogenesis trajectories was defined by its absence of neuroblasts and instead relied on pro-neuro ependymoglia intermediates. Interestingly, glutamatergic neuron cluster 6, which showed transcriptional similarity to turtle aDC and mouse cortex, was part of this trajectory. It will be exciting to investigate the lineage potential of ependymal cells and neuroblasts in the axolotl using methods such as genetic barcoding that have allowed us to gain a better understanding of the lineage relationships of radial glia, progenitor cells and neurons in the mammalian forebrain (56).

In contrast to mammalian glial cells, salamander ependymoglia show neurogenic activity after injury. We found that axolotl ependymoglia go through an injury-specific transcriptional state defined by the upregulation of genes involved in wound healing and cell migration. Interestingly, early upregulated genes *Runx1* and *Kazald1* have also been shown to be expressed in the mesenchymal limb blastema of axolotl and Xenopus laevis (57, 58). *Runx1* is also expressed in planarian neoblasts after amputation (59) suggesting that these genes are involved in a generic injury response regardless of cell type and species. Apart from the early wound response, regenerative neurogenesis is strikingly similar to homeostatic neurogenesis. Astonishingly, we find that *Satb1+*/*Rorb+* glutamatergic neurons are regenerated after injury and re-establish input connectivity from the olfactory bulbs. The conservation of transcriptional profiles, connectivity and potential function of these neurons makes them ideally suited targets to study recovery of functional neuronal circuits.

## ACKNOWLEDGEMENTS

We would like to thank the Treutlein, Camp and Tanaka labs for their discussions, input, and support. FACS sorting support was provided by the single-cell facility at D-BSSE, ETH Zurich. Illumina sequencing was done in the Genomics Facility at DBSSE, ETH Zurich. We thank the IMP Bio-Optics facility and the animal caretaker team (Vienna Biocenter) for outstanding service.

## Author contributions

KL, AM, TG, EMT, BT designed the study. KL prepared and collected the samples, preformed lineage tracing, and staining. AM Performed single-nuclei experiments. TG, AM, JSF analyzed the sequencing data. All authors prepared the figures, wrote and approved the manuscript.

## Data and materials

Code is available at GitHub 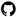 (https://github.com/tomasgomes/pallium_evo).

## Materials and Methods

### Axolotl strains and maintenance

White (*d/d*) axolotls not biased to a specific sex were used for all experiments. All lines were bred and maintained in IMP facilities and each animal is kept individually. All handling and surgical procedures were carried out in accordance with the local ethics committee guidelines. Animal experiments were performed as approved by the Magistrate of Vienna (Genetically Modified Organism Office and MA58, City of Vienna, Austria, license GZ51072/2019/16 and license GZ665226/2019/21). Caggs:LoxP-eGFP-3polyA-LoxP-Cherry (Caggs:lp-Cherry) transgenic animals were described previously (61). Animals were anesthetized in 0.03% benzocaine (Sigma-Aldrich, E1501) before electroporation or surgery. Axolotl husbandry was performed as described previously (61). 10-11cm animals were used for all sequencing experiments.

### Nuclei isolation

Frozen pallium microdissections and whole pallium dissections were dissociated for generating single-nuclei gene-expression libraries following a modified protocol from 10x (Demonstrated protocol CG000365, Rev B). In brief, we prepared and precooled wash and lysis buffers before moving tissue pieces to a precooled 1.5mL tube. 50µL of lysis buffer was added to the sample and dissociated via 2-5 short pulses with an electric grinder. The pestle of the grinder was washed with a 150µl wash buffer before centrifugation for 5 min at 500xg (4°C). Supernatant was removed and the pellet gently washed with 200µl of wash buffer before centrifugation for 5 min at 500xg (4°C). Supernatant was removed and the pellet was resuspended in 50-100µl of PBS + 0.5% BSA. Resulting nuclei were then assessed (count and viability) using Trypan Blue assay, counted using the automated cell counter Countess (Thermo Fisher).

### Brain injury

Brain injuries were performed as described previously (8). Animals were deeply anesthetized in 0.03% benzocaine and rectangular cranial skin/skull flaps above the right telencephalon hemisphere were performed using scalpels, forceps and spring scissors. A 1mm × 1mm piece of dorsolateral telencephalon was removed and the injury was always positioned between the choroid plexus and the olfactory bulb. After the injury, the cranial skin/skull flaps were restored without suture and the axolotls were returned to individual tanks.

### EdU administration

Anesthetized axolotls were injected intraperitoneally with 800 µM EdU (diluted in 1xPBS) at a dosage of 10 µl/g. FastGreen dye (Sigma-Aldrich) was added to the injection mix to aid visualization. Injected axolotls were kept out of water for a 15 min recovery period under benzocaine-soaked towels. After recovery, injected axolotls were returned to water. Following the desired pulse-chase period, axolotls were sacrificed and brains harvested.

### Nuclei isolation for Div-Seq

Frozen pallium dissections were dissociated for generating single-nuclei gene-expression libraries similarly to described above with some modifications (44). In brief, we prepared and precooled wash and Div-seq lysis buffers before moving tissue pieces to a precooled 1.5mL tube. 50µL of lysis buffer was added to the sample and dissociated via 2-5 short pulses with an electric grinder. The pestle of the grinder was washed with a 150µl wash buffer before centrifugation for 5 min at 500xg (4°C). Supernatant was removed and the pellet gently washed with 200l of wash buffer before centrifugation for 5 min at 500xg (4°C). Supernatant was removed and the pellet was resuspended in 55µl of wash buffer. Resulting nuclei were then assessed (count and viability) using Trypan Blue assay, counted using the automated cell counter Countess (Thermo Fisher).

EdU staining was performed immediately using Click-iT EdU Flow Cytometry assay Kit (Thermo Fisher Scientific, #C10424), 500 µl reaction buffer was added directly to the resuspension buffer (mix is made following the manufacturer’s protocol), mixed well and left in RT for 30min. 3ml of wash buffer was added to the resuspended nuclei and mixed well, then nuclei were spun down for 5 min at 500xg (4°C), supernatant was removed and nuclei were resuspended in 500 µl PBS + 0.5% BSA with DAPI and FACS sorted immediately.

### Preparation of single-nucleus RNA and ATAC profiles

For snRNAseq profiling, nuclei were diluted to an appropriate concentration to obtain approximately 2,000-10,000 nuclei per lane of a 10x microfluidic chip device. Single-nuclei cDNA was synthesized per manufacturer recommendations (10x Genomics v3.1) before continuing to library preparation with 25% of the total cDNA volume. Combined snRNAseq and snATACseq were generated with the Chromium Single Cell Multiome ATAC + Gene Expression kit following manufacture recommendations (10x Genomics). Final libraries were sequenced on Illumina NovaSeq SP or S1 flow cell.

### Spatial transcriptomics with Visium

Whole axolotl brain was flash frozen before it was embedded in a prechilled optimal cutting temperature compound. The sample was then set into a dry ice bath with isopentane until frozen and stored at 80°C. Cryosections were cut at a thickness of 10µm, adhered to Visium (10x) spatial transcriptomic (ST) slides (10x) and stored at 80°C until the following day. Tissue slices were fixed in cold methanol before being stained with hematoxylin and eosin. ST slides were imaged as recommended on a Nikon T2i at 20x using a tile scan over all slice sections. Following image capture, tissue slices were permeabilized. Optimal permeabilization conditions were determined by using the Tissue Optimization kit (10x), and the optimal time was found to be 52 min. Spot-captured RNA was reverse transcribed before second-strand synthesis and cDNA denaturation. qPCR was used to determine the optimal number of cDNA amplification cycles as recommended by the manufacturer. cDNA was amplified using 18 cycles before continuing to Visium spatial gene expression library construction. Visium libraries were sequenced on the Illumina NovaSeq SP following sequencing recommendations.

### Gene expression quantification

UMI counts for single-nuclei and Visium data were obtained with kallisto v0.46.2 (62) and bustools v0.41.0 (63). Quantification was based on the Ambystoma mexicanum transcriptome v4.7, based on the genome v6.0 release (64). To account for an increased number of reads of intronic origin due to nuclei isolation, the full exonic and intronic regions were considered for standard gene expression analyses. Expression values were summarized to gene names according to the transcriptome annotation. For each transcript, a human transcriptome-base annotation was preferred, otherwise the annotation obtained from all other species was used. If neither were present, the axolotl gene ID was used as the gene name. Following quantification, detection of nuclei-containing droplets was performed by using the intersection of the results from the EmptyDrops and DefaultDrops function in the DropletUtils package (65), before saving the data as a Seurat object (66). For Visium data, an additional step used SpaceRanger to retrieve the image spot coordinates. These quantification workflows were implemented using Nextflow (67).

Spliced and unspliced gene expression counts for all datasets used in RNA velocity analysis were obtained using a pipeline similar to the one used for general quantification. Additionally, a new kallisto transcriptome index was produced, which explicitly included differentiated exons and introns plus overhangs, and a step was added to differentially obtain the spliced and unspliced fractions using bustools’ “capture” function.

The spatial matching of Visium spots to the tissue was done using SpaceRanger from which only the mapping information was extracted and no quantification was performed.

### Open chromatin detection and quantification

The snATACseq fraction from the multiome sequencing samples was aligned and quantified using CellRanger-ARC. This requires simultaneous quantification of RNA and ATAC, yet only the latter was kept, together with the RNA quantification using kallisto-bustools. Quantification for each sample used 36 threads and 130Gb RAM and took approximately 1.5 days.

Due to limitations on contig size for the creation of STAR (68) genome index files, the axolotl genome (v6.0) was first processed into contigs smaller than 500Mb, while avoiding breaking annotated genes (+/-10kb) into separate contigs. Indexing the genome took one day using 98Gb and 32 threads.

To allow for downstream DNA-based analysis, a “BS.genome” R package (https://bioconductor.org/packages/release/bioc/html/BSgenome.html) was built for the axolotl genome processed into 500Mb contigs.

### snRNAseq data quality control and integration

Standard procedures for filtering, variable gene selection, dimensionality reduction, and clustering were performed using the Seurat (v3.1, (66)) in RStudio using R. Cells with fewer than 200 genes and mitochondrial transcript proportion higher than 40% were excluded. We used DoubletFinder (69) to identify and exclude doublet cells. Counts were log-normalized. Data was then scaled for the top 10,000 variable genes, while regressing out the number of reads and percent mitocondria.

Gene expression per cell was then projected using Principal Component Analysis (PCA). The top 20 PCs were selected based on inspection of an elbow plot of variance explained, and used to perform a Uniform Manifold Approximation and Projection (UMAP) for visualization. Harmony (70) was used to integrate data of different 10x chemistries (v3.1 and multiome v1, fig. S1, B to C).

### snRNAseq iterative clustering, differential gene expression, and annotation

Identification of brain cell types followed an iterative clustering approach. First, 20 PCs were used to generate a nearest-neighbors graph. Clustering was then performed using the Louvain algorithm and a resolution of 0.7. Cells were then divided into neuronal and non-neuronal based on combining expression patterns of canonical cell type markers, e.g. *Gli2, Aqp4, Snap25, Slc17a6/7, Gad1, Gad2, Pdgfra, Ptprc*, and *Cd34*.

Annotation of specific cell types was followed, relying on iterative subsetting and processing, where the top 10,000 variable genes were selected and scaled per cell, and the top PCs were selected to perform UMAP for visualization and build a nearest-neighbors graph for Louvain clustering. Clusters were then annotated based on known marker genes for cell types.

The subset of non-neuronal cells used 20 PCs and a resolution of 0.9 for Louvain clustering. The canonical markers used for annotation in this group included *Gli2, Aqp4, Pdgfra, Ptprc*, and *Cd34*. From these, ependymoglia cells were subset. Dimensionality reduction relying on 25 PCs and clustering with a resolution of 1.1 revealed 15 distinct clusters. GO term enrichment (fig. S5, B to C) was calculated using the gProfiler2 R package (71).

Neuronal cells were subset, and 30 PCs were used. Clusters were further identified at a resolution of 1.1. Cells were then annotated as either GABAergic, Glutamatergic, or neuroblast based on expression of *Gad1/2, Slc17a6/7*, or *Mex3a* respectively. GABAergic (*Gad1/2+*) neuronal cells were processed using the top 30 PCs and a Louvain resolution of 1.1, which revealed 30 clusters. Glutamatergic (*Slc17a6/7+*) neuronal cells were processed using the top 30 PCs and a Louvain resolution of 1.1, which revealed 29 clusters. Due to a lower number of features and few differentially expressed genes, cluster 23 was deemed representing low-quality cells and removed from further analysis. Neuroblasts (*Mex3a+*) were processed using the top 25 PCs, and clustering at a resolution of 1.7 revealed 15 clusters.

Differentially expressed genes among clusters of each subset (ependymoglia, GABAergic, Glutamatergic, and neuroblast) were identified using the wilcoxauc method from presto (72) with ‘pseudocount.use’ set to 0.1 (table S1). The average expression for the top 200 DE genes for each cluster was used in the hierarchical clustering of all populations (fig. S1F).

### Cross-species cell and gene comparisons

Axolotl brain cell types were compared with cells sampled from mouse (3, 38, 39) and turtle (2) brain. Pairwise ortholog gene correspondences were directly obtained from Ensembl (73, 74). Spearman correlation was used for pairwise cross-species comparisons, using the corr.test function in the psych R package. Correlation was calculated on the normalized average expression for each cluster considered. The genes were adjusted to the scope of each comparison by using the intersection of pairwise DE genes between all pairs of clusters considered for each species, and used all those genes of just those coding for transcription factors where mentioned.

### Pallium region prediction for whole pallium snRNA-seq

Pallium microdissected snRNA-seq gene expression data was used to predict the original dissected region. Using scikit-learn (75), a probability-calibrated Logistic Regression Classifier was trained on 80% of the data, stratified by identified cell type and region. On the test set, the model had an F1 score of 0.997 and an accuracy per cell type no lower than 85%.

The trained model was used to predict the region in the snRNA-seq fraction from the whole pallium multiome data. Additionally, cell type region of origin was taken into account when deciding the final region assignments, since cell types were annotated in an integrated version of whole and microdissected data. In cell types where more than 30% of cells were assigned a maximum probability lower than 90% of belonging to a given region, the cells not passing that threshold were assigned as “Undetermined” region. This is because lower assignment probabilities were in general more prevalent in cell types originating from more than one brain microdissected region.

### Mapping cell types into Visium spatial data

Cell types were mapped to Visium data using cell2location (76). The different chemistries (v3.1 and multiome) were used as batch variables, and the region and animal were considered covariates in the regression model. For the spatial mapping, 25 cells were assumed per location.

To compare NB progenitor brain region mapping, the 0-1 scaled average score of each spot was taken for VGLUT+ (Glut 0, 1, 2, 3, 4, 7, 9, 11, 13) and GABA+ (5, 6, 8, 10, 12) NBs, and their difference was plotted (fig. S5J).

### snATACseq quality control and processing

Initial analysis of the snATAC-seq fraction of axolotl pallium multiome data was performed using Signac (77). Peaks detected by CellRanger-ARC were used, for a total of 631,645. Cells were kept if they had a number of reads or features between 200 and 10,000, a nucleosome signal lower than 2, a TSS enrichment of at least 3, and no more than 5% of reads from mitochondrial origin.

### RNA velocity and pseudotime analysis

RNA velocity analysis was performed using the python packages scvelo (41) and CellRank (42). Glutamatergic cell types were assigned into 5 groups according to their similarities with the neuroblast progenitors (fig. S7A). All trajectories included all activated ependymoglia cells (clusters 3 and 4 - active ependymoglia). Glutamatergic neuron cell types that did not conform to a trajectory were excluded from the analysis.

For the UMAP dimensionality reduction, the top 10000 variable genes were used, and 15 neighbors were used for the moments calculation using 15 PCs. CellRank was then used to determine the assignment probabilities to the end state cell types as well as the ependymal starting state, using the top 150 cells according to the scvelo inferred latent time and end states.

Within each group, a global pseudotime was obtained by multiplying the maximum probability of each cell being assigned any glutamatergic fate with the inverse probability of assigning the ependymoglia fate. This ensures that all cells derive from the root population, and diverge according to their calculated probabilities.

Genes were determined to be differentially variable along pseudotime by fitting a Generalized Additive Model modeling the interaction between each fate and the pseudotime (modeled as a natural spline with 5 degrees of freedom), weighted by the fate probabilities. Lineage specificity genes were also obtained using the compute_lineage_drivers function from CellRank. Genes were then determined to be lineage-specific within a specific group if they significantly differed between lineages along pseudotime (adjusted p-value<=0.05), and were either uniquely detected as lineage drivers by CellRank for that lineage alone or had a difference greater than 20% in expression compared to the other lineage. The correlation values resulting from compute_lineage_drivers were compared for the same trajectories obtained with steady-state and pallium regeneration data (fig. 6M, fig. S8, H to K).

### GRN construction from multiome data

We used the R package Pando (43) to infer a GRN from multiome data of cells from all Glutamatergic trajectories. To enrich potential regulatory regions, peaks were linked to genes with correlated expression using the Signac function LinkPeaks() with a distance threshold of 10 mb. All peaks with significant (p < 0.05) linkage to genes were used as candidate regions for Pando (initiate_grn(), exclude_exons=FALSE). To find transcription factor binding sites in candidate regions, we obtained vertebrate motif annotations from the CORE collection of JASPAR2020 (75) and used them as the input for the Pando function find_motifs(). Next, we inferred the GRN using the Pando function infer_grn() considering a 10Mb region upstream and downstream of the TSS. We used a gaussian generalized linear model (model=‘glm’) to infer regulatory coefficients from log-normalized transcript counts and binarized peak counts summarized to louvain clusters. We corrected for multiple testing using the Benjamini-Hochberg method to obtain an FDR-adjusted p-value, to which a significance threshold of 0.05 was applied.

### A. Visualization of the GRN

All transcription factors in the network were visualized based on both co-expression and regulatory relationships. First, we computed the Pearson correlation between log-normalized expression of all transcription factors in the network. From the correlation value r and estimated model coefficient β between all transcription factors i and j, we then computed a combined score s as: 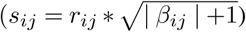 resulting in a TF by TF matrix. We performed PCA on this matrix and used top 20 PCs as an input for UMAP as implemented in the uwot R package (https://github.com/jlmelville/uwot) with default parameters.

To highlight subgraphs for individual TFs, we computed the shortest path from the TF to every gene in the GRN graph. Next, we filtered the paths by only retaining the path with the lowest average log10 p-value for each target gene.

### Differential accessibility of regulatory regions

To constructy trajectory-specific subgraphs of the GRN, we tested regulatory regions for differential accessibility between neuronal trajectories. For this, we fit a generalized linear model with binomial noise and logit link for each peak i on binarized peak counts Y with the total number of fragments per cell and the trajectory label as the independent variables: *Y*_*i*_ ∼ *n*_*fragements* + *trajectory*_*label*. In addition, we fit a null model, where the trajectory label was omitted: *Y*_*i*_ ∼ *n*_*fragments*. We then used a likelihood ratio test to compare the goodness of fit of the two models using the lmtest R package (version 0.9) (https://cran.r-project.org/web/packages/lmtest/index.html). Multiple testing correction was performed using the Benjamini-Hochberg method. We applied a threshold of 0.05 on the FDR-corrected p-value to identify trajectory-specific regulatory regions. Trajectory-specific GRNs were constructed by pruning all non-specific edges in the global GRN.

### Div-seq quality control and processing

Standard procedures for filtering, variable gene selection, dimensionality reduction, and clustering were performed using the Seurat (v3.1) in RStudio using R. Cells with fewer than 200 genes and mitochondrial transcript proportion higher than 40% were excluded. We used DoubletFinder (69) to identify and exclude doublet cells. Counts were log-normalized. Data was then scaled for the top 10,000 variable genes, while regressing out the number of reads and percent mitocondria.

Gene expression per cell was then projected using Principal Component Analysis (PCA). The top 30 PCs were selected based on inspection of an elbow plot of variance explained. Harmony was used to integrate data of different experimental batches (batch 1-4). The resulting integrated projection was used to perform a Uniform Manifold Approximation and Projection (UMAP) for visualization. Clustering was then performed using the Louvain algorithm and a resolution of 1.3, which were used to estimate the variation of cell type proportions along regeneration (fig. 6D).

### Div-seq region and cell type automatic annotation

Cells from the Div-seq dataset were classified into cell types and brain regions like the steady-state data by training two models, using both the snRNA-seq and multiome gene expression fraction.

To predict the brain region, a probability-calibrated Logistic Regression Classifier was trained on 80% of the data, stratified by identified cell type and region. For cells from the multiome fraction, the predicted region was used, and cells with ‘Undetermined’ regions were excluded. The resulting model had an F1 score of 0.955 on the test set.

To predict the cell type, a probability-calibrated Random Forest Classifier was trained on 80% of the data, stratified by identified cell type and region. The resulting model had an F1 score of 0.820 on the test set.

As a complementary visualization, we mapped Glutamatergic neurons (fig. 6J), as well as GABAergic neurons and neuroblasts (fig. S8, E to F) to the UMAP projections obtained in our previous steady-state analyses. Div-seq nuclei were mapped at the UMAP coordinates of the steady-state cell with the highest Spearman correlation coefficient.

### Integration of ependymal cells from different protocols

Ependymoglia were collected from the steady-state and Div-seq datasets. Cells were jointly normalized, and the top 10,000 genes were used for downstream analysis. Data was scaled and number of UMI counts, percent of mitochondrial reads, and the dataset of origin were regressed out. 15 PCs were selected from the elbow plot, and used for integration by protocol (v3.1, multiome) and Div-seq batch (1-4) with Harmony, with a tau parameter of 30. A subsequent UMAP was generated with the top 30 components from Harmony.

Clustering was done using the Louvain algorithm with a resolution parameter of 1.5. Marker genes were obtained using the presto R package (table S4). GO term enrichment was calculated using the gProfiler2 R package (71). GO terms were then grouped into 10 clusters by semantic similarity using the GOSemSim package (79) to simplify interpretation. From these, illustrative terms were selected to group genes (fig. 6H and table S4).

### Div-seq RNA velocity analysis

RNA velocity analysis for the Div-seq dataset was performed similarly to the uninjured dataset.

Groups of neuroblasts and differentiated neurons were selected to be identical to the groups in the steady-state neurogenesis. This is supported by the very similar neurolast-glutamatergic neurons observed in the injured and uninjured datasets (fig. S7A and fig. S8G).

For the UMAP dimensionality reduction, the top 10000 variable genes were used, and 50 neighbors were used for the moments calculation using 25 PCs. CellRank was then used to determine the assignment probabilities to the end state cell types as well as the ependymal starting state, using the top a random set of 50 nuclei out of the top 150 according to the scvelo inferred latent time and end states. This change in the number happens due to the lower number of nuclei in the end states of the trajectory.

Within each group, a global pseudotime was obtained by multiplying the maximum probability of each cell being assigned any glutamatergic fate with the inverse probability of assigning the ependymoglia fate. This ensures that all cells derive from the root population, and diverge according to their calculated probabilities.

### Brain electroporation

Intraventricular injections of the seCre plasmid (diluted in 1x PBS at concentrations varying from 0.5 µg/µl to 2.5 µg/µl, mixed with FastGreen dye (Sigma-Aldrich) to aid visualization) was performed on anesthetized animals. The skin on top of the brain was removed using a scalpel and tweezers and a glass microneedle was inserted into the ventricle of the midbrain. The plasmid solution was injected until the ventricles of the forebrain were completely filled with solution. Animals were covered with a Whatman paper soaked in 1xPBS and electroporations were performed with a NEPA21 electroporator (Nepagene) using tweezers with round platinum plate electrodes 2 or 5 mm in diameter depending on the size of the animals (2mm electrodes for 4cm animals, 5 mm electrodes for 10cm animals). Unidirectional (+) and bidirectional (+/-) pulses were used. One 70V pore-forming pulse for 5 ms was followed by four 30V transfer pulses for 50ms each with a pulse interval of 999 ms and with a decay of 10% between the pulses.

### Neurobiotin injection

Axolotls were fully anesthetized and 10% Neurobiotin diluted in water and FastGreen dye for visualization was pressure injected into the desired region of the brain. Brains were harvested 24 hours after injection.

### Brain tissue collection for HCR or immunohistochemistry

Axolotls were fully anesthetized, decapitated with scissors and brains were extracted and fixed overnight at 4°C in 4% paraformaldehyde. Fixed brains were washed 6 times 30 minutes each with 1xPBS and either incubated overnight in 30% sucrose in 1xPBS at 4°C if used for cryosectioning or used immediately for wholemount immunohistochemistry.

### HCR probe design

cDNA sequences for genes of interest were first analyzed for unique regions. Probe pairs were designed using a custom made python script. oPool of oligos at 50pmol containing up to 37 probe pairs in a pool for each gene were ordered from IDT. Detailed sequences of each probe can be found in Supplementary Table 3.

### Cryosections

Cryoprotected brains were embedded in optimal cutting temperature (OTC) compound, frozen in liquid nitrogen and stored at −80°C until sectioning. Coronal cryosections of 18 µm thickness were prepared from frozen blocks and stored at −20°C until use.

### HCR in situ hybridization on cryosections

HCR in situ hybridization was performed according to the manufacturer’s instructions by Molecular Instruments with the following modifications: Probe concentration was increased to 0.8 pmol and slides were covered with parafilm for all incubation steps. Slides were mounted in a Glycerol mounting medium (80% glycerol, 1xPBS, 20mM Tris pH 8, 2.5 mg/mL propyl gallate).

### Immunohistochemistry on cryosections

Cryosections were rehydrated in 1xPBST (0,2% TritonX-100) for 30 min at room temperature. In case antigen retrieval was needed slides were heated to 90°C in a Tris-based buffer (Vector Laboratories, H-3301) The slides were allowed to cool for at least 30 min and washed in 1xPBST before incubation with primary antibodies The respective primary antibodies were diluted in 1% NGS overnight at 4°C. After 6 washes for 30 minutes each at room temperature the secondary antibodies were applied in 1% NGS together with DAPI (1:500 dilution in 1xPBST of 5 mg/ml stock) for 3 hours at 37°C. Slides were mounted with 60% glycerol and stored at 4°C until imaging.

### Wholemount immunohistochemistry and Ethylcinnamate clearing of brains

Fixed and washed brains were incubated in ice cold acetone at −20 degrees for 15 minutes and washed with 1xPBST (0,2% TritonX-100) 3 times for 30 minutes. Blocking was performed using 4% Goat Serum, 1% DMSO in 1xPBST for 1 hour at room temperature on a horizontal shaker. Primary antibodies were diluted in 4% Goat Serum, 1% DMSO in 1xPBST and incubated over three nights at 4°C on a horizontal shaker. Wash 6 times 30 min with 1xPBST at room temperature on a horizontal shaker. Secondary antibodies were diluted in 4% Goat Serum, 1% DMSO in 1xPBST, incubated over three nights at 4°C on a horizontal shaker. Finally, brains were washed 6 times 30 min with 1xPBST at room temperature on a horizontal shaker.

Clearing of Cre-lox tracing brains was performed using an Ethylcinnamate-based clearing approach (80). First, brains were dehydrated in a sequential series of 1-Propanol in 1xPBS (30%, 50%, 70%, 100%, 100%) which was pH-adjusted to 9-9.5. Each incubation step was performed for 9-15 hours. Refractive index matching was performed using Ethylcinnamate. Brains were incubated for at least 1 hour room temperature and used for imaging when fully transparent.

### Wholemount staining and CUBIC clearing of Neurobiotin-injected brains

Staining and clearing of Neurobiotin-injected brains was performed using an CUBIC-based clearing approach using CUBIC-L, CUBIC-R1a and CUBIC-R+(N) solutions ((81) and http://www.cubic.riken.jp). Fixed and washed brains were incubated in 50% CUBIC-L/R1a solution (mixed 1:1 and diluted to 50% in dH2O) for 3 hours on a shaker at room temperature. Afterwards brains were incubated with 100% CUBIC-L/R1a solution for 30 mins on a shaker at 37°C, followed by washes with 1xPBS for 6 times 10 min each at room temperature. Anti-Streptavidin was used at a dilution of 1:1000 4% Goat Serum, 1% DMSO in 1xPBST and incubated over three nights at 4°C on a horizontal shaker. Brains were washed 6 times 30 min with 1xPBST at room temperature on a horizontal shaker and finally incubated in CUBIC refractive index matching solution CUBIC-R+(N), on a horizontal shaker until cleared.

### Microscopy

Cleared brains were mounted in glass bottom dishes (Ibidi) and imaged on an inverted Zeiss LSM980 Axio Observer (inverted) confocal microscope with a 10x/0.3 EC plan-neofluar objective. Cryosections were imaged on an inverted Zeiss LSM980 Axio Observer (inverted) confocal microscope using a 20x/0.8 plan-apochromat objective. ZenBlue 3.2 was used for image acquisition and automatic stitching. Image preparation was performed using FIJI (based on ImageJ 1.53c).

## Supplementary Data Figures

**fig. S1.**
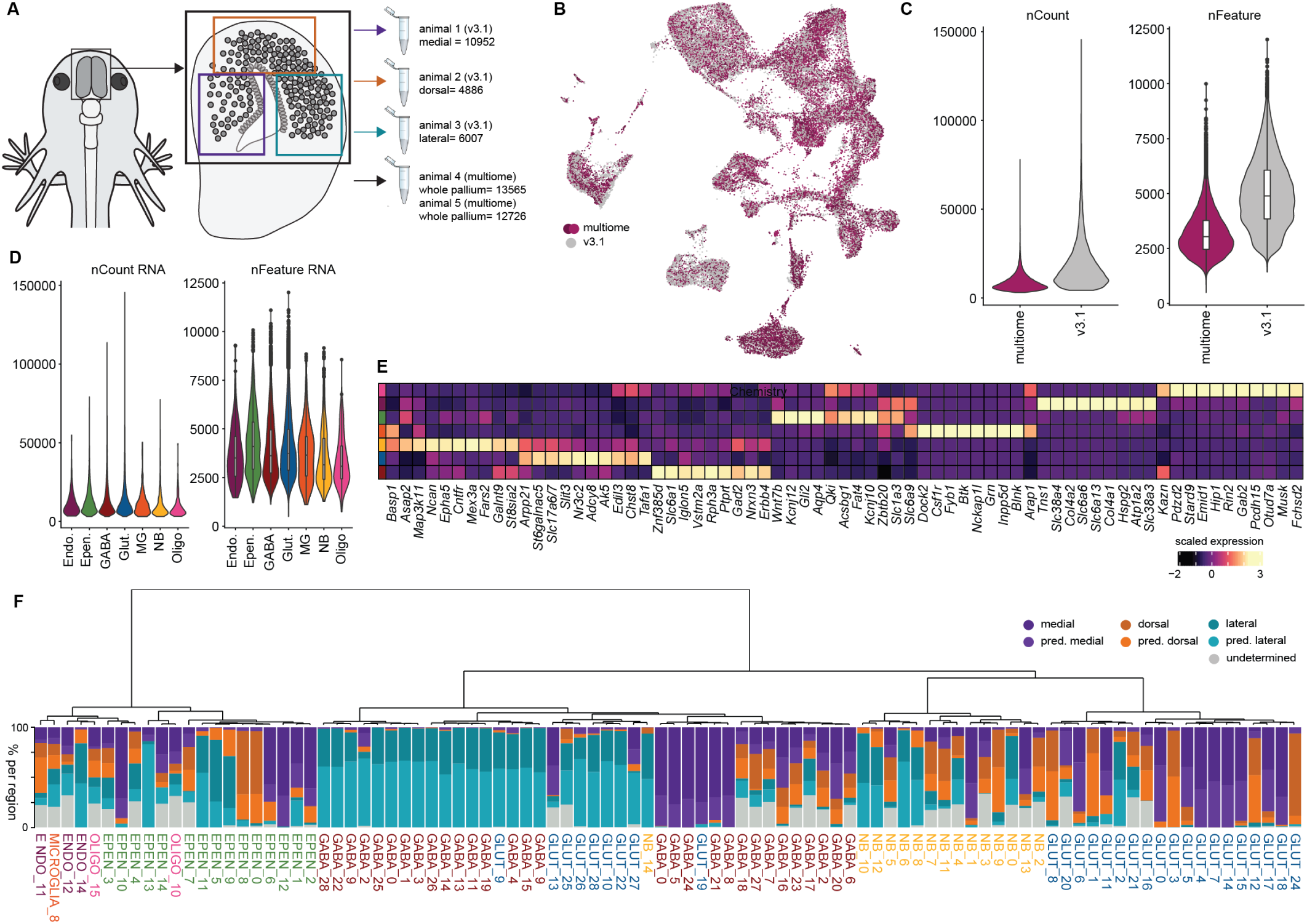
Global experimental overview of the axolotl pallium single-nuclei sequencing. (A) Schematic highlighting the regions of the axolotl telencephalon and chemistry used for single nucleus RNA sequencing, as well as nuclei number for each sample.(B) UMAP of all nuclei colored by chemistry.(C) Violin plots of counts and features grouped by chemistry. (D) Violin plots of counts and features grouped by cell type. (E) Heatmap of top marker genes for each cell type. (F) Dendrogram based on the mean expression of the top 200 differentially expressed genes for each cell cluster. Stacked barplot illustrating the regional distribution of the populations of cells. GABA, GABAergic neuron; Glut, Glutamatergic neuron; NB, neuroblast; Epen, ependymoglia cell; Endo., Endothelial cell, MGs, microglia; Oligo., oligodendrocyte.

**fig. S2.**
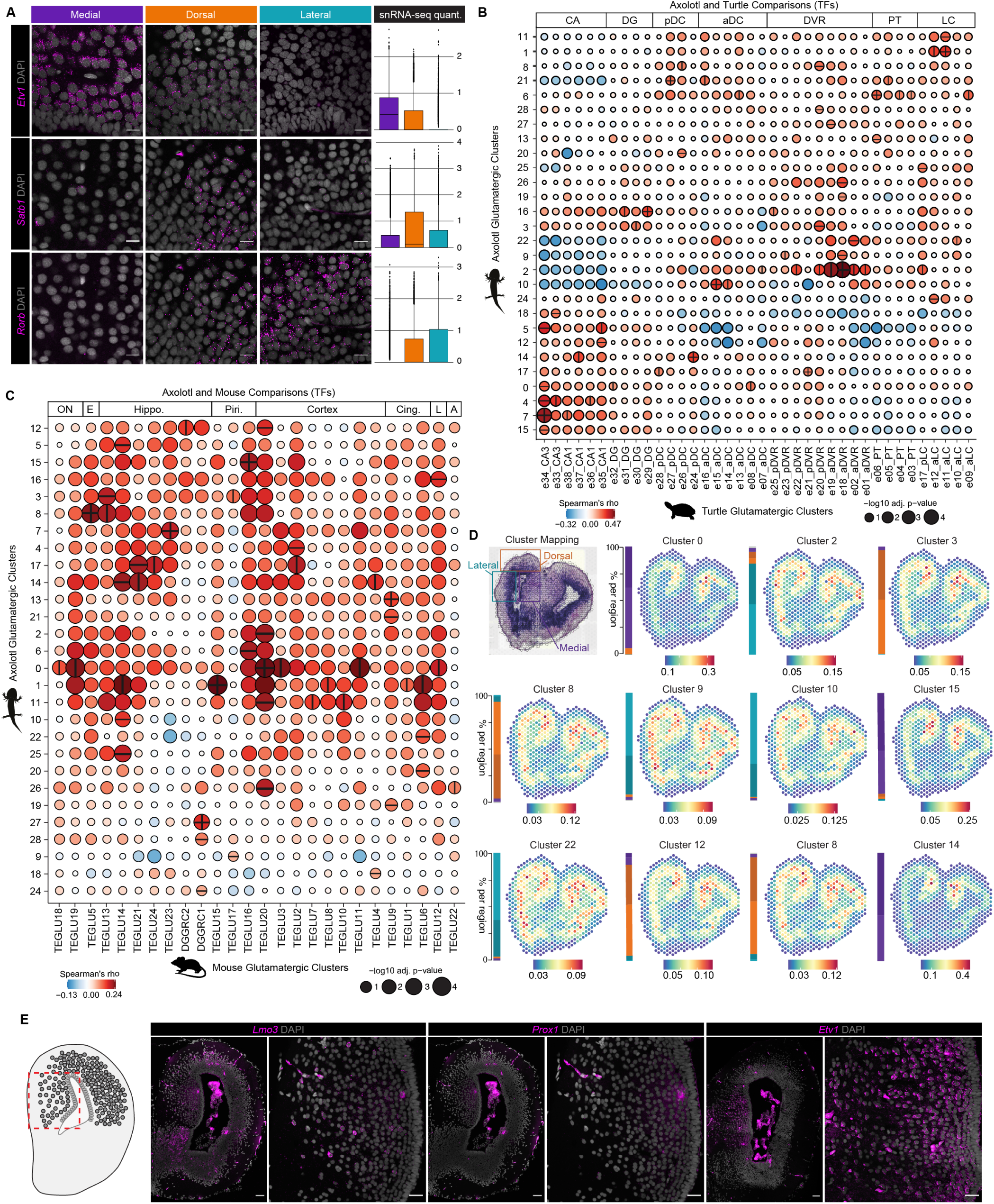
Additional characterization of Glutamatergic neuron populations. (A) HCR in situ hybridizations and snRNAseq quantifications for *Etv1, Satb1*, and *Rorb* in medial, dorsal and lateral regions. Scale bars are 25 µm. (B) Correlation analysis between transcription factor expression of axolotl glutamatergic neuron types and turtle glutamatergic neuron types (data from (2)). (C) Correlation analysis between transcription factor expression of axolotl glutamatergic neuron types and mouse glutamatergic neuron types (data from (3)). TEGLU, Telencephalon projecting excitatory neurons. DGGRC, Dentate gyrus granule neurons. (D) H&E stained Visium slice, annotated by region. Spatial Mapping of select glutamatergic neuron clusters. Stacked barplot illustrating the regional distribution of the populations of cells. (E) Schematic of whole pallium slice, red dashed box indicates medial pallium region. HCR in situ hybridizations and snRNAseq quantifications for *Lmo3, Prox1*, and *Etv1* in whole pallium and medial pallium regions. Scale bars are 25 µm.

**fig. S3.**
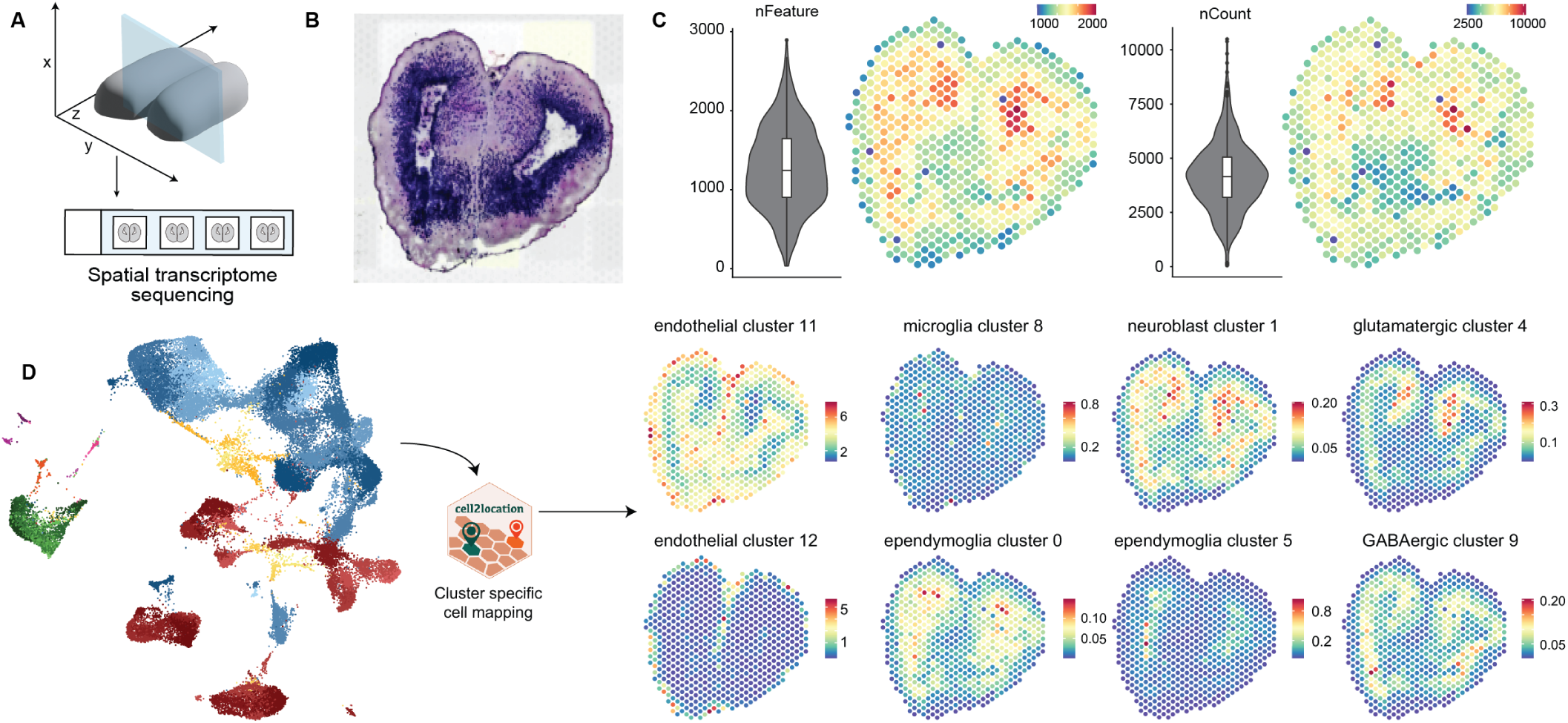
Technical overview of axolotl pallium spatial transcriptomics data. (A) Schematic tissue slice for Visium spatial transcriptomics. (B) H&E stained Visium slice. (C) Violin plots and spatial mapping of counts and features across Visium spots. (D) Schematic of spatial mapping approach using cell2location (60) and examples of cell cluster mapping.

**fig. S4.**
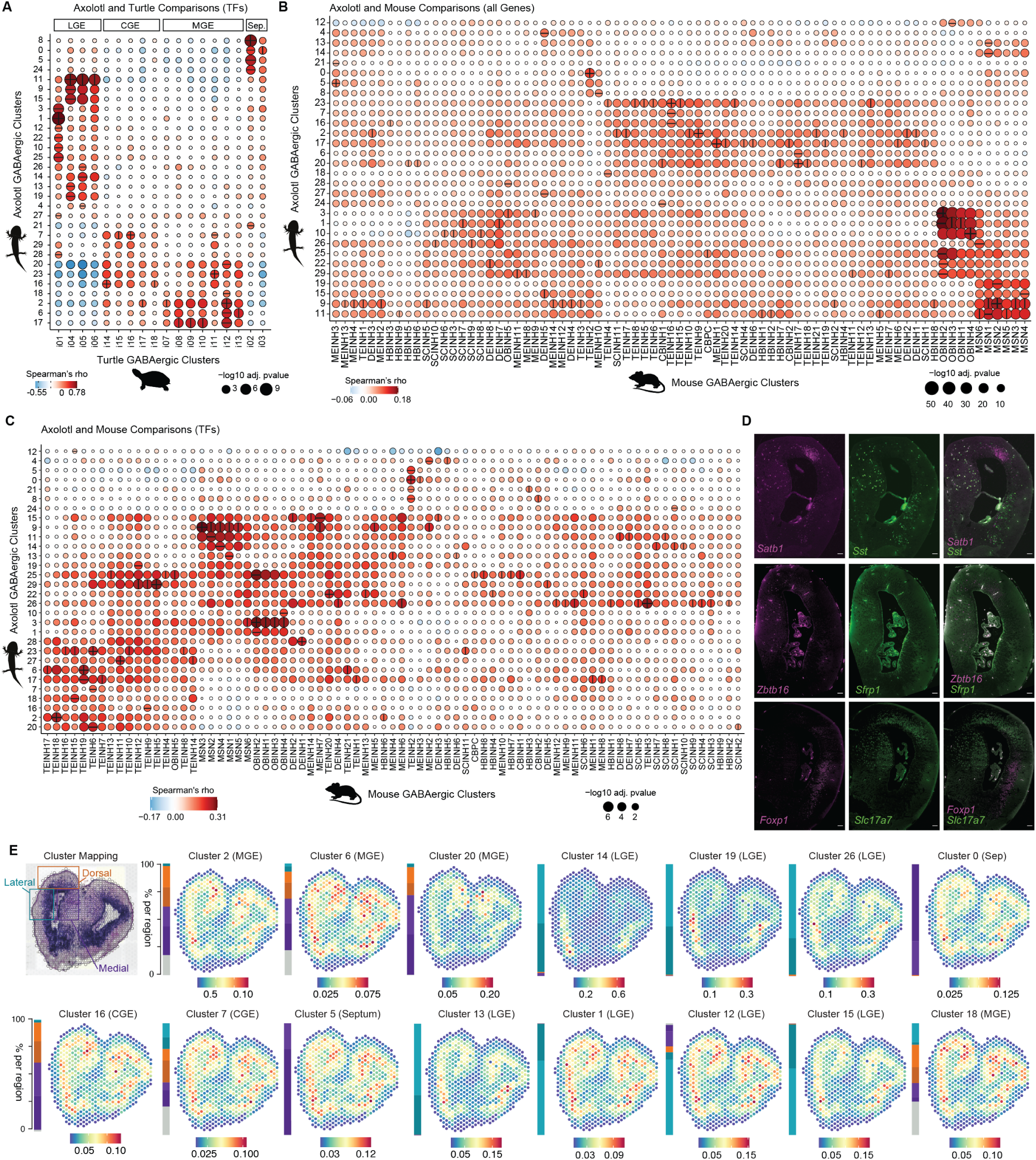
Additional characterization of GABAergic neuron populations. (A) Correlation analysis between transcription factor expression of axolotl GABAergic neuron types and turtle GABAergic neuron types (data from (2)). (B) Correlation analysis between gene expression profiles of axolotl GABAergic neuron types and mouse GABAergic neuron types (data from (3)). MEINH, Di- and mesencephalon inhibitory neurons. TEINH, Telencephalon inhibitory interneurons. DEINH, Di- and mesencephalon inhibitory neurons. HBINH, Hindbrain neurons. SCINH, Spinal cord inhibitory neurons. OBINH, Olfactory inhibitory neurons. CBPC, Cerebellum neurons. MSN, Telencephalon projecting inhibitory neurons. (C) Correlation analysis between transcription factor expression of axolotl GABAergic neuron types and mouse GABAergic neuron types (data from (3)). MEINH, Di- and mesencephalon inhibitory neurons. TEINH, Telencephalon inhibitory interneurons. DEINH, Di- and mesencephalon inhibitory neurons. HBINH, Hindbrain neurons. SCINH, Spinal cord inhibitory neurons. OBINH, Olfactory inhibitory neurons. CBPC, Cerebellum neurons. MSN, Telencephalon projecting inhibitory neurons. (D) HCR in situ hybridizations for *Satb1, Sst, Zbtb16, Sfrp1, Foxp1*, and *Slc17a7*. Scale bars are 100 µm. (E) H&E stained Visium slice, annotated by region. Spatial mapping of select glutamatergic neuron clusters. Stacked barplot illustrating the regional distribution of the populations of cells.

**fig. S5.**
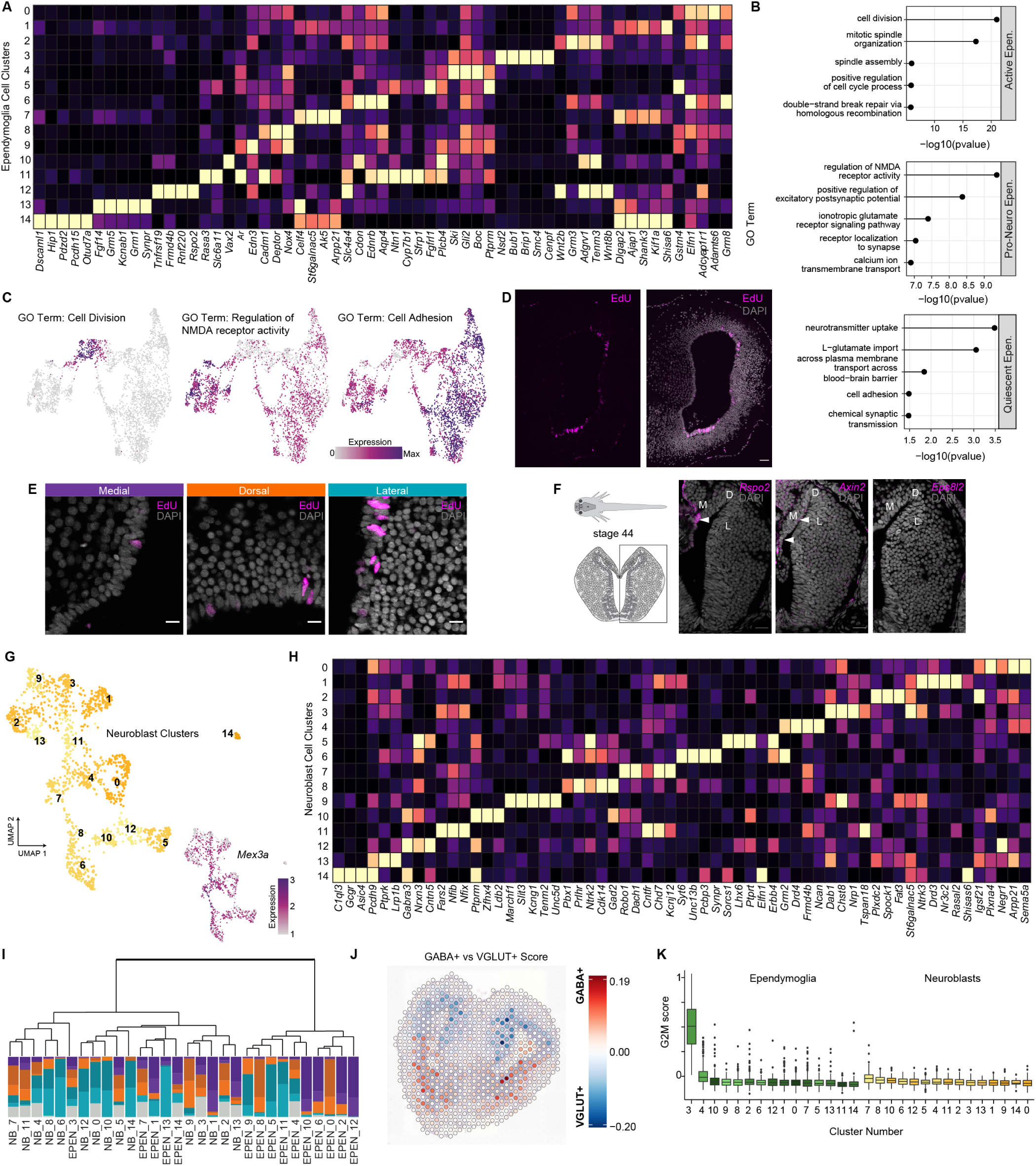
Additional characterization of Ependymal cells and Neuroblasts. (A) Heatmap of top marker gene expression across 15 ependymoglia clusters. (B) Top GO terms for each class of ependymoglia (active, pro-neuro, and quiescent). (C) UMAP plot of ependymoglia clusters colored by GO term expression. (D) EdU only staining (left) and EdU with DAPI staining (right). (E) EdU staining after 2 consecutive injections within 2 weeks and snRNAseq qualifications for G2M and S phase score in medial, dorsal and lateral regions.. Scale bars are 25 µm. (F) Schematic of stage 44 axolotl and corresponding brain slice. HCR in situ hybridizations *Rspo2, Axin2*, and *Eps8l2* in the stage 44 developing pallium. Scale bars are 50 µm. (G) UMAP plots colored by neuroblast clusters (top) and *Mex3a* gene expression (bottom). (H) Heatmap of top marker gene expression across 15 neuroblast clusters. (I) Dendrogram clustered by top 200 differentially expressed genes. (J) Visium spots colored by GABA+ (red) and VGLUT+ (blue) scores. (K) Boxplot of G2M score for ependymoglia and neuroblast clusters.

**fig. S6.**
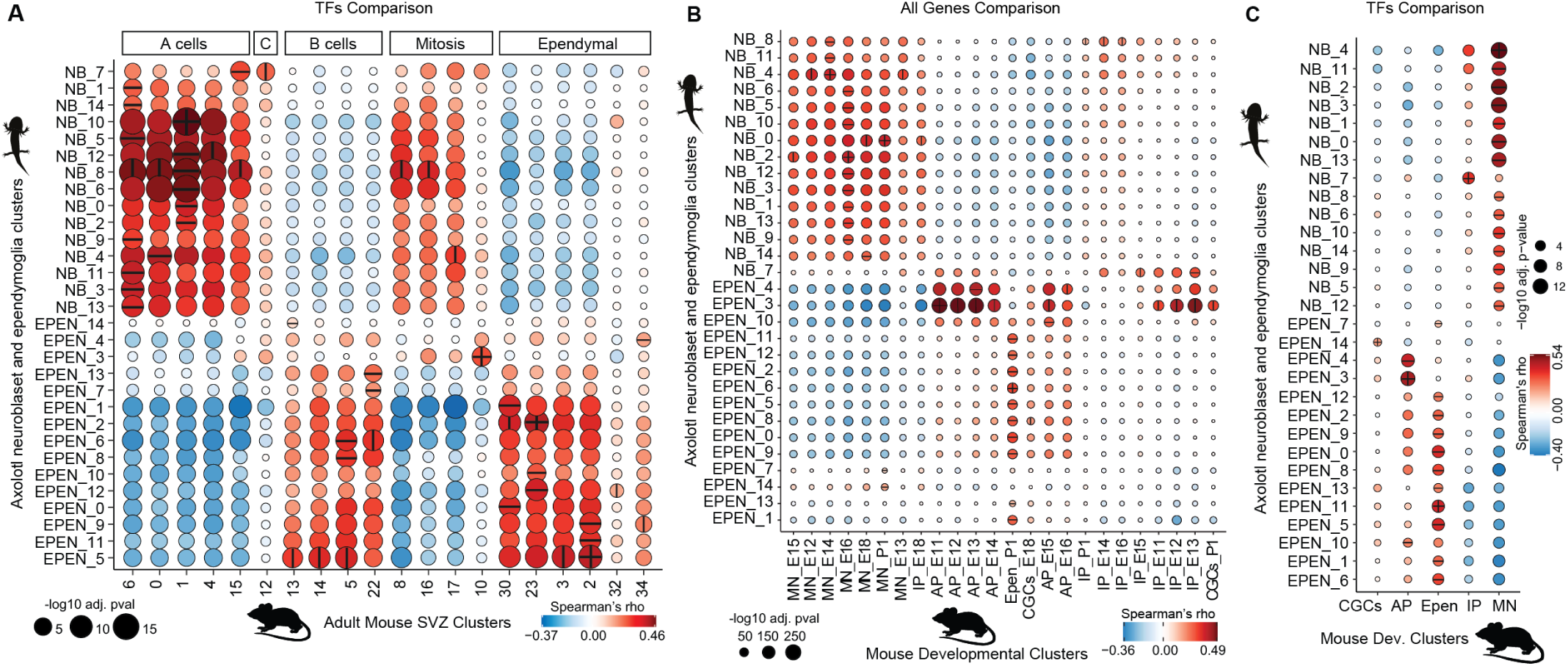
Complementary cross-species comparisons of Ependymal cells and Neuroblasts. (A) Correlation analysis between transcription factor expression of axolotl neuroblasts, ependymoglia and adult mouse SVZ cell types (data from (38)). (B) Correlation analysis between expression profiles of axolotl neuroblasts, ependymoglia and developing mouse cell types (data from (39)). MN, migrating neuron. IP, intermediate progenitor. AP, apical progenitor. Epen, ependymal cell. CGC, cycling glial cells. (C) Correlation analysis between transcription factor expression of axolotl neuroblasts, ependymoglia and developing mouse cell types (data from (39)). MN, migrating neuron. IP, intermediate progenitor. AP, apical progenitor. Epen, ependymal cell. CGC, cycling glial cells.

**fig. S7.**
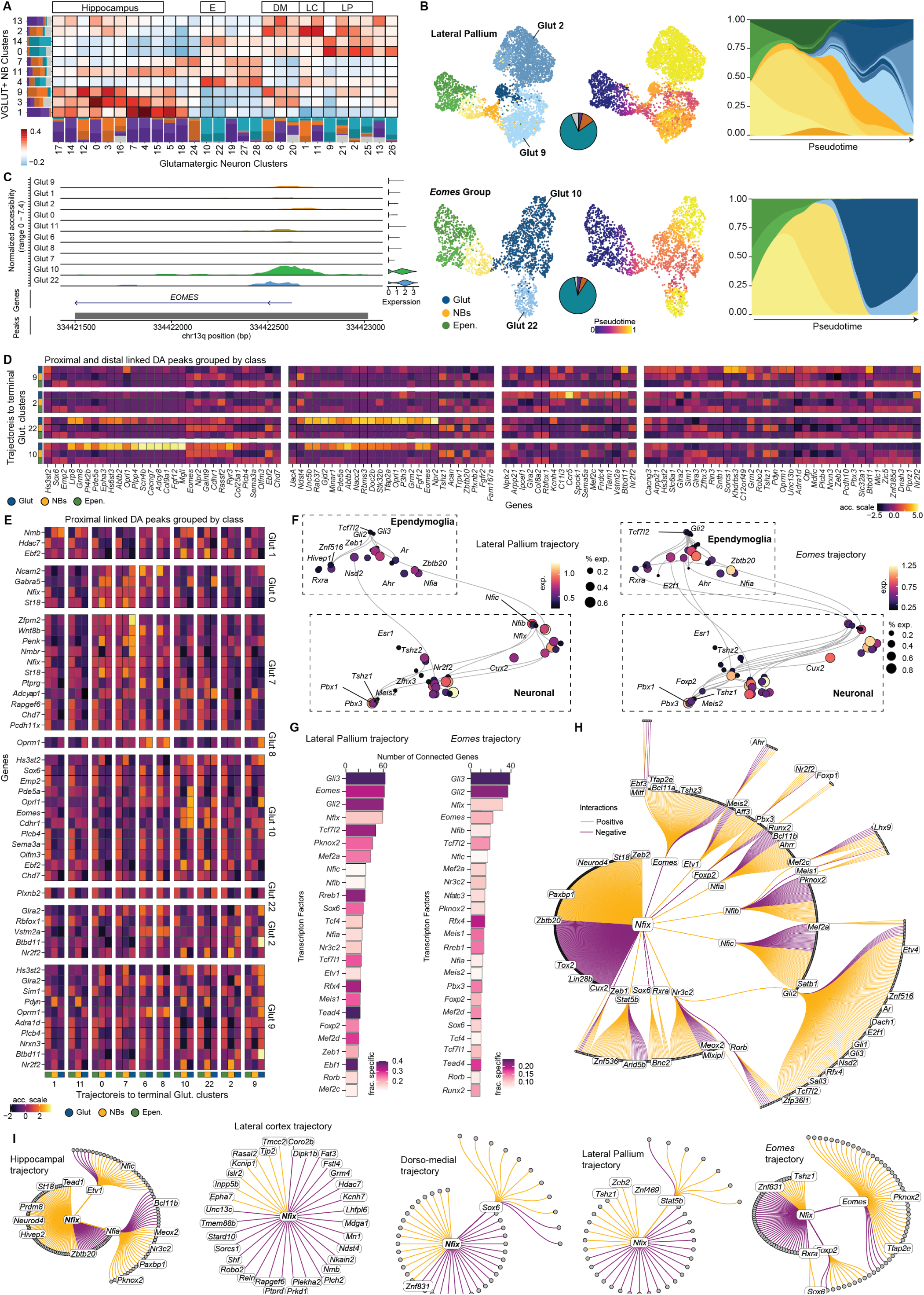
Additional information and trajectories of axolotl pallium steady-state neurogenesis. (A) Heatmap of transcriptional similarity between glutamatergic neuron and neuroblast clusters. Stacked barplot illustrating the regional distribution of the populations of cells. Box labels indicate the groups of cells with greatest similarity and thus used for RNA velocity trajectories. E, *Eomes*. DM, Dorsal-medial. LC, Lateral cortex. LP, Lateral pallium. (B) Glutamatergic trajectories reflecting adult neurogenesis of lateral pallium and Eomes groups from fig. S7A. UMAPs colored by cell types (left) and pseudotime (right). Pie charts represent the regional composition of neuron clusters. Pseudotemporal cell type progression from ependymoglia to glutamatergic neurons during neurogenesis. (C) Representative example peaks associated with *Eomes* for all terminal glutamatergic clusters identified from RNA velocity analysis (Fig. 5B and fig. S7A). (D) Heatmap of chromatin accessibility changes in distal and proximal elements for glutamatergic clusters 10, 22, 2, and 9. (E) Heatmap of chromatin accessibility changes in proximal elements for glutamatergic clusters 1, 0, 7, 8, 10, 22, 2, and 9. (F) Trimmed GRN UMAP embedding of the inferred gene modules based on co-expression and inferred interaction strength between transcription factors for lateral pallium (left) and *Eomes* (right) trajectories. Color scale indicates expression and size represents the percent of cells expressing. (G) Barplot of the top 25 transcription factors ranked by number of connections for each transcription factor for lateral pallium (left) and Eomes (right) trajectories. (H) Global regulatory network centered on *Nfix*. Yellow indicates positive regulation of *Nfix* and purple indicates negative regulation of *Nfix*. (I) *Nfix* regulatory networks specific for each trajectory. Yellow indicates positive regulation of *Nfix* and purple indicates negative regulation of *Nfix*.

**fig. S8.**
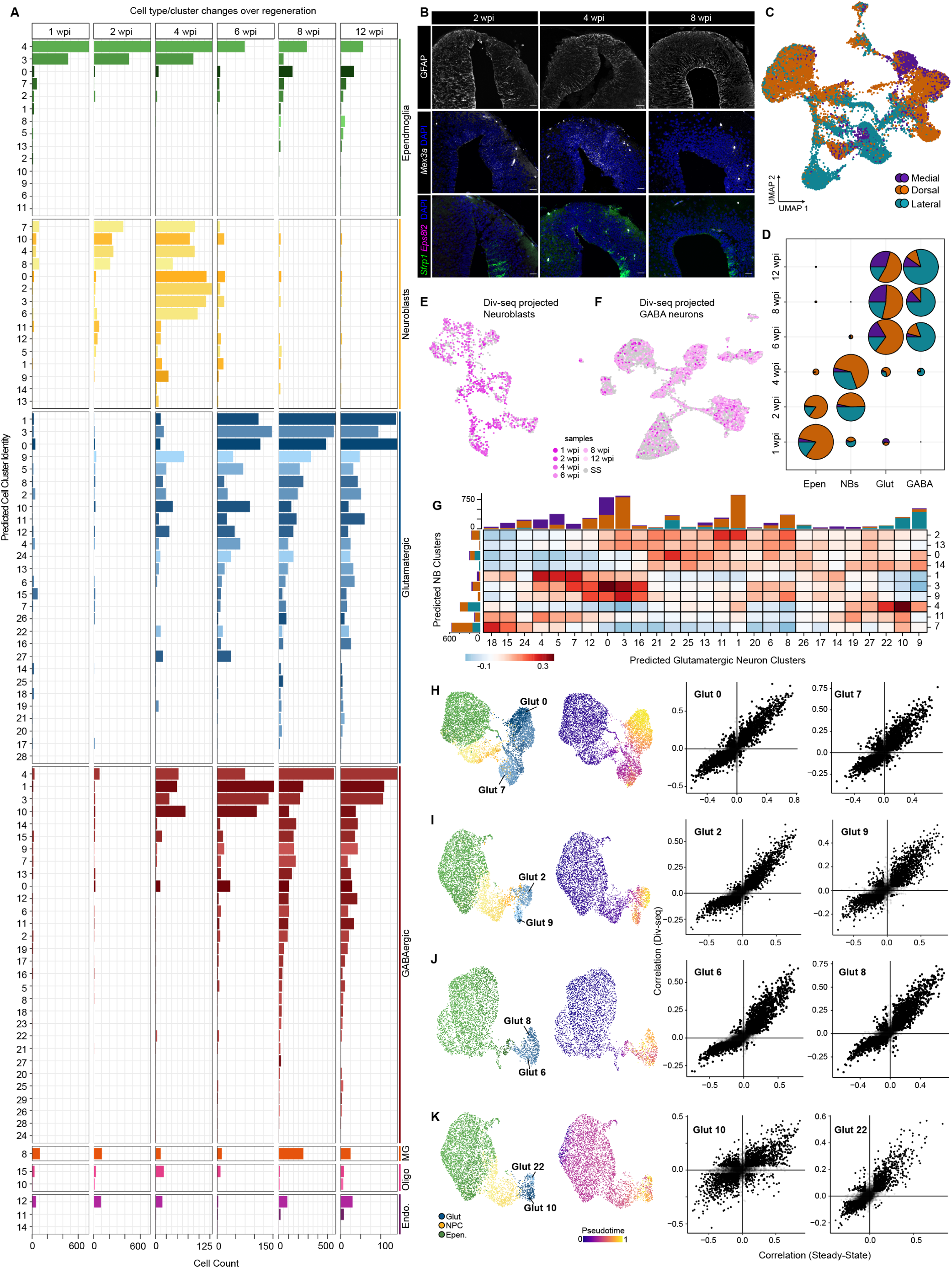
Extended characterization of axolotl pallium regeneration. (A) Barplot of number of nuclei captured for each predicted cell cluster from steady state from each Div-seq time point. (B) HCR in situ hybridizations and antibody staining for GFAP, *Mex3a, Eps8l2*, and *Sfrp1* at 2, 4, and 8 wpi. Scale bars are 100 µm. (C) UMAP of all Div-seq nuclei colored by region prediction. (D) Pie charts colored by predicted region and grouped by predicted cell type and Div-seq time point.(E) Correlation projection of all Div-Seq neuroblasts (pink) to steady state neuroblasts (gray). (F) Correlation projection of all Div-Seq GABAergic neurons (pink) to steady state GABAergic neurons (gray). (G) Heatmap of transcriptional similarity between predicted glutamatergic neuron and predicted neuroblast clusters. Barplots illustrate the predicted regional distribution and amount of Div-seq nuclei. (H) Trajectories reflecting regenerative neurogenesis of glutamatergic neuron clusters 0 and 7. UMAP colored by cell type (left) and pseudotime (right). Correlations of lineage driver genes with the assignment probability for Glut 0 (left) and Glut 7 (right) trajectories, in regenerative from Div-Seq (y-axis) and to Steady-state (x-axis) neurogenesis. (I) Trajectories reflecting regenerative neurogenesis of glutamatergic neuron clusters 2 and 9. UMAP colored by cell type (left) and pseudotime (right). Correlations of lineage driver genes with the assignment probability for Glut 2 (left) and Glut 9 (right) trajectories, in regenerative from Div-Seq (y-axis) and to Steady-state (x-axis) neurogenesis. (J) Trajectories reflecting regenerative neurogenesis of glutamatergic neuron clusters 6 and 8. UMAP colored by cell type (left) and pseudotime (right). Correlations of lineage driver genes with the assignment probability for Glut 6 (left) and Glut 8 (right) trajectories, in regenerative from Div-Seq (y-axis) and to Steady-state (x-axis) neurogenesis. (K) Trajectories reflecting regenerative neurogenesis of glutamatergic neuron clusters 10 and 22. UMAP colored by cell type (left) and pseudotime (right). Correlations of lineage driver genes with the assignment probability for Glut 10 (left) and Glut 22 (right) trajectories, in regenerative from Div-Seq (y-axis) and to Steady-state (x-axis) neurogenesis.

